# Generation of Systemic Chimeras via Rabbit Induced Pluripotent Stem Cells Reprogrammed with KLF2, ERAS, and PRMT6

**DOI:** 10.1101/2024.01.10.575048

**Authors:** Florence Perold, Hong-Thu Pham, Yannicke Pijoff, Nathalie Doerflinger, Sylvie Rival-Gervier, Anaïs Moulin, Luc Jouneau, Bertrand Pain, Thierry Joly, Véronique Duranthon, Marielle Afanassieff, Pierre Savatier, Nathalie Beaujean

## Abstract

Little is known about the molecular underpinnings of pluripotent stem cells’ (PSCs) ability to colonize the epiblast of preimplantation embryos and generate chimeras. In our study, using rabbit PSCs as a model system, we conducted unbiased screening of a cDNA library that encodes a panel of 36 pluripotency factors. From this screening, we identified KLF2, ERAS and PRMT6, whose overexpression confers the ability for self-renewal in a KOSR/FGF2-free culture medium supplemented with LIF, activin A, PKC and WNT inhibitors. The reprogrammed cells acquired transcriptomic and epigenetic features of naive pluripotency, including the reactivation of the 2^nd^ X-chromosome. Leveraging these PSC lines, we determined the transcriptomic signature of embryonic colonization-competence, demonstrating transcriptional repression of genes involved in MAPK, WNT, HIPPO, and EPH signaling pathways, alongside the activation of genes involved in amino-acid metabolism, NF-kB signaling, and p53 pathway. Remarkably, a subset of reprogrammed cells, expressing CD75 at a high level, gained the ability to produce chimeric fetuses with a high contribution from PSCs in all lineages.

## Introduction

Systemic chimerism, often designated germline chimerism in rodents, is the ability of naïve pluripotent stem cells (PSCs) to contribute to the formation of all tissues and organs within both the fetus and the adult organism after injection into blastocysts or aggregation with morulae. To date, systemic chimerism has been documented exclusively in rodents, and it is seen as the definitive test for pluripotency, providing incontrovertible evidence that the injected cells can generate fully functional cells in every organ and tissue. However, there currently exists no equivalent experimental demonstration of pluripotency in non-rodent species, including lagomorphs and non-human primates (NHPs). The pluripotency of rabbit and monkey PSCs– whether embryo-derived PSCs or induced PSCs (iPSCs)–is presently inferred from *in vitro* differentiation and experimental teratomas, which are not comprehensive tests. The production of systemic chimeras in rabbits and NHPs represents a significant and challenging goal. Chen and colleagues made groundbreaking contribution by reporting the first–and to date, the only– two NHP fetuses with widespread PSC colonization in cynomolgus monkeys (Chen et al. 2015). Interestingly, they detected sporadic PSC-derived VASA^+^ cells in the testis, which suggests germline colonization. However, the rate of chimerism was acknowledged to be low and highly variable between tissues and organs. In rabbits, Tapponnier and colleagues reported two fetuses in which less than one cell in 1000 was derived from injected iPSCs (Tapponnier et al. 2017). This suggests the need for further research into the chimera formation process to understand its limitations and find potential solutions.

Studies conducted in mice showed that only PSCs in the naïve state of pluripotency can colonize preimplantation embryos and contribute to germ layer differentiation. PSCs that self-renew in the primed state of pluripotency, known as EpiSCs (Brons et al. 2007; Tesar et al. 2007), cannot contribute to chimeras in the same experimental setting, unless they are genetically engineered to express a high level of E-cadherin or BCL2 (Nichols and Smith 2009) (Ohtsuka et al. 2012; Masaki et al. 2016). This clear association between the naïve state and preimplantation embryo colonization observed in rodents seems far less distinct in primate species, including human and NHPs. While some studies reported the formation of chimeric fetuses after injecting naïve PSCs from humans or macaque monkeys into mouse and macaque embryos (Gafni et al. 2013; Fang et al. 2014; Chen et al. 2015; Cao et al. 2023), other studies observed very low, or no colonization in the examined embryos and fetuses (mouse, macaque and pig) (Masaki et al. 2015; Theunissen et al. 2016; Honda et al. 2017; Fu et al. 2020). Many critical factors differ between these studies, such as cell lines, reprograming protocols, and host species, complicating any definitive conclusion. However, when examining the early fate of naïve PSCs after injection into host embryos, human and NHP PSCs appear less capable than mouse PSCs of surviving, multiplying, and colonizing the epiblast (Masaki et al. 2015; Aksoy et al. 2021; Tan et al. 2021). This is unless they are engineered to overexpress the polycomb factor BMI1 (Huang et al. 2018), the anti-apoptotic factor BCL2 (Aksoy et al. 2021; Roodgar et al. 2022), or inactivate TP53 (Bayerl et al. 2021). Aksoy and colleagues showed that naïve human and NHP PSCs prematurely differentiate when injected into rabbit morulae, while mouse ESCs actively proliferate and colonize the host epiblast (Aksoy et al. 2021). Altogether, these findings strongly suggest that most human and NHP PSC lines are intrinsically unfit for producing chimeric organisms with strong organ colonization.

Given these findings, it is crucial to decipher the molecular underpinnings of PSCs’ ability to effectively colonize preimplantation embryos. However, addressing this issue in NHPs presents substantial challenges due to the intrinsic difficulties related to embryo production and their limited availability. Given these hurdles, we propose the use of rabbits as a beneficial intermediary model. Rabbits provide two primary benefits: firstly, rabbit embryos are readily available in large quantities, and secondly, PSCs from both primates and rabbits share many similarities, especially regarding their growth requirements (Osteil et al. 2013; Afanassieff et al. 2015; Osteil et al. 2016; Savatier et al. 2017). Leveraging rabbit iPSCs and embryos as our model system, we conducted unbiased screening of a cDNA library encoding a panel of 36 pluripotency factors. From this, we identified genes whose overexpression endows the ability to produce chimeric embryos and fetuses with high contribution from PSCs in all major organs.

## Results

### Development of a novel culture regimen and reporter cell line for naïve pluripotency in rabbit

Rabbit iPS cells (rbiPSCs) are commonly cultured in basal medium supplemented with FGF2 and knockout serum replacement (KOSR) factors, or KF medium (Osteil et al. 2013; Tapponnier et al. 2017). When attempting to grow rbiPSCs (B19 cell line (Osteil et al. 2013)) in media originally described for naive human and rhesus monkey PSCs (*i.e*. HENSM (Bayerl et al. 2021), 5iLA (Theunissen et al. 2014), t2iLGöY (Takashima et al. 2014), 4i/L/b (Fang et al. 2014), and LCDM (Yang et al. 2017b)), the rbiPSCs either died or differentiated. Stable self-renewal was not observed (**Fig. S1A**). However, when cultured in murine embryonic fibroblast (MEF)-conditioned N2B27 basal media supplemented with 10 ng/ml activin, 10,000 U/ml LIF, 250µM Vitamin C, 2.5µM protein kinase C inhibitor Gö6983, and 2.5µM Tankyrase inhibitor XAV939 (hereafter called VALGöX), B19_KF cells displayed progressive morphological and molecular changes. After 48h in VALGöX (referred to as B19_VAL), they exhibited an increased nuclear-cytoplasmic ratio and formed more compact colonies (**Fig. 1A, 1B**). Immunofluorescence analysis also revealed higher expression of two naïve markers, DPPA5 and OOEP (Bouchereau et al. 2022), increased levels of permissive histone mark H3K14ac, and decreased levels of repressive histone mark H3K9me3, H3K27me3, and 5-methyl-Cytosine (5mC), which suggests the onset of a primed to naïve-like pluripotency transition (**Fig. 1B**).

**Figure 1:**
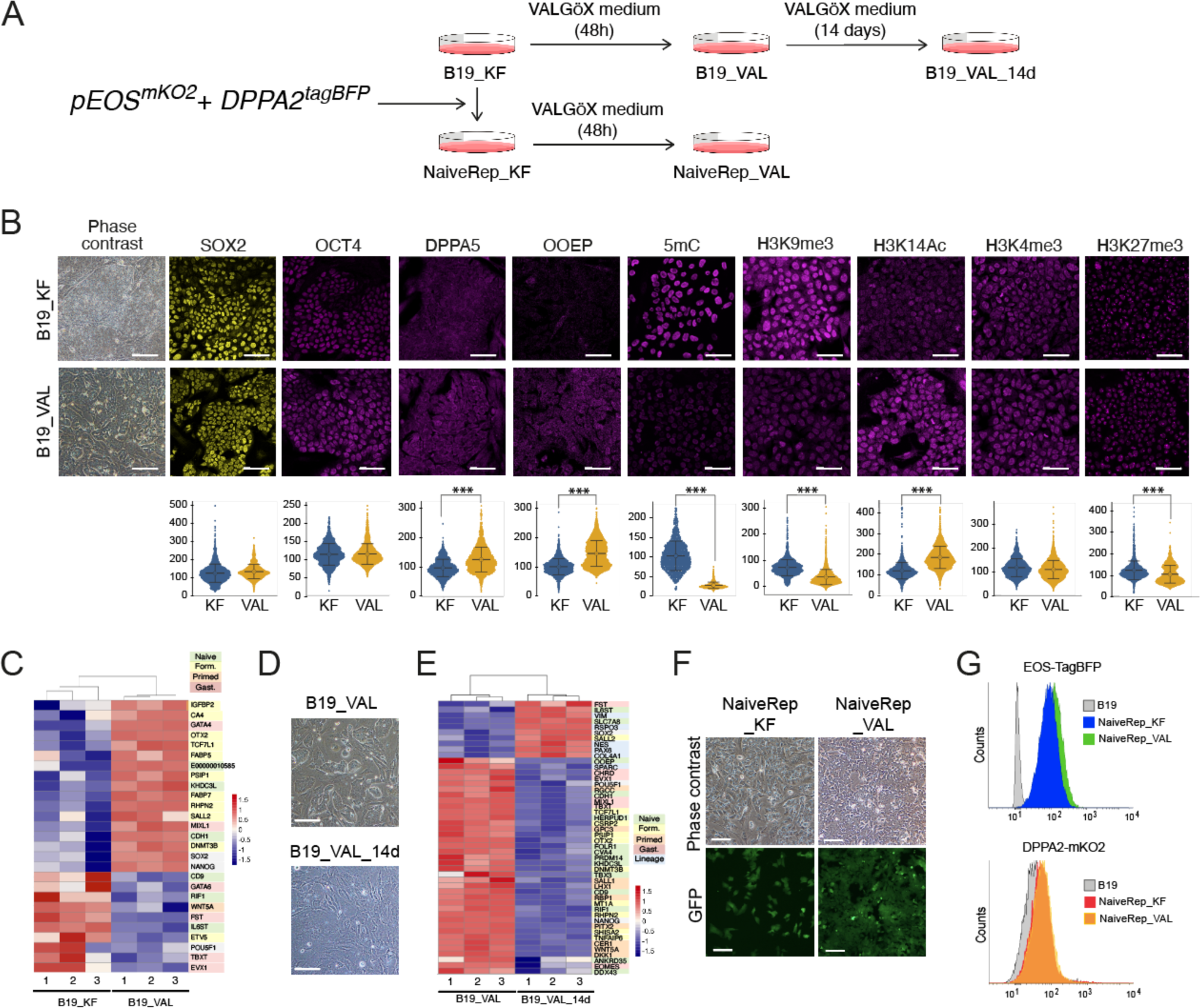
Development of a novel culture regimen and reporter cell line for naïve pluripotency in rabbit. (**A**) Experimental scheme of B19_VAL, B19_VAL_14d, NaiveRep_KF and NaiveRep_VAL cell line generation. (**B**) Immunostaining of SOX2, OCT4, DPPA5, OOEP, 5mC, H3K9me3, H3K14Ac, H3K4me3, and H3K27me3 in B19 cells before (_KF) and after (_VAL) switching to VALGöX culture medium. Violin plots represent the distribution of fluorescence intensities. Welch’s unequal variances t-test was performed to compare KF and VAL conditions. A difference is considered significative (***) if difference between sample means is greater than 10% and the p-value less than 1e-10. Scale bars: 50 µm for immunostainings, 100µm for phase contrast. **C**) Heatmap representation of differentially expressed genes between B19_KF and B19_VAL cells. (**D**) Phase contrast images of B19 cells 48h (_VAL) and 14 days (_VAL_14d) after switching to VALGöX culture medium. Scale bars: 200 µm. (**E**) Heatmap representation of differentially expressed genes between B19_VAL and B19_VAL_14d cells. (**F**) Phase contrast and epifluorescence images of NaiveRep_KF and NaiveRep_VAL cells. Scale bars: 100 µm. (**G**) Flow cytometry analysis of parental B19_KF (control iPSC), NaiveRep_KF and NaiveRep_VAL showing blue and red fluorescence associated with EOS-tagBFP and DDPA2-mKO2, respectively.

A transcriptome analysis of B19 cells performed before (B19_KF) and 48 hours after switching to the VALGöX culture regimen showed reduced expression of rabbit primed and gastrulation markers, and increased expression of rabbit formative markers (Bouchereau et al. 2022) (**Fig. 1C**). However, B19_VAL cells lacked certain key characteristics of naïve PSCs. Specifically, immunostaining for H2AK119ub and H3K27me3, two histone post-translational modifications associated with X chromosome inactivation in mice and humans (Chaumeil et al. 2011), exhibited nuclear foci in 80% (H2AK119ub) and 78% (H3K27me3) of B19_VAL cells, and in 69% (H2AK119ub) and 75% (H3K27me3) of B19_KF cells (**Figs. S1B, S1C**). This suggests the presence of an inactive X chromosome, typical of primed PSCs, in the majority of both B19_KF and B19_VAL cells. We also assessed the colonization ability of B19_KF and B19_VAL cells after injection into rabbit preimplantation embryos. B19_KF and B19_VAL cells express a green fluorescent protein (GFP) under the transcriptional control of the CAG ubiquitous promotor (Osteil et al. 2013; Tapponnier et al. 2017). Eight cells were injected into early morula stage embryos (E2.8), and the morulae were cultured for three days, developing into late blastocysts (∼E5). Both B19_VAL and B19_KF cells demonstrated limited capacity for epiblast and trophectoderm lineage contribution (**Fig. S1D**). In summary, these findings suggest that the VALGöX culture regimen promotes the development of rabbit iPSCs towards a more immature pluripotent state, but does not achieve genuine naïve-state reprogramming. After 14 days in VALGöX, B19_VAL cells displayed morphological differentiation (**Fig. 1D**). Transcriptomic analysis revealed a decline in the expression of naïve, formative and primed pluripotency markers, alongside an up-regulation of lineage markers confirming the initiation of differentiation (**Fig. 1E**). The VALGöX culture regimen therefore fails to support B19 cell self-renewal.

In order to monitor the transition from primed to naïve pluripotent states and identify rare cells undergoing this conversion in B19 cell cultures, we developed a dual fluorescent naïve pluripotency reporter. *DPPA2* was selected as a naïve pluripotency marker in rabbits based on its expression profile in pre-implantation embryos. Single-cell RNAseq analyses showed that *DPPA2* is expressed in all pluripotent cells of day 3 (early-blastocyst stage) and day 4 (mid-blastocyst stage) rabbit embryos, with expression decreasing from day 5 onwards, suggesting down-regulation during the naive to primed pluripotency transition (**Fig. S1E**) (Bouchereau et al., 2022). B19 cells expressing the GFP were transfected with Dox-inducible Cas9D10A and gRNAs targeting the rabbit *DPPA2* locus, as well as a plasmid carrying monomeric Kusabira Orange (*mKO2*) with *DPPA2* homology sequences. The homology-directed repair strategy aimed to integrate mKO2 downstream of the ATG codon of *DPPA2* (**Fig. S1F**). After G418 selection and genomic PCR analysis, one cell clone was chosen and further transfected with *pEOS^tagBFP^*, a reporter plasmid containing a blue fluorescent protein (tagBFP) under the naive-specific distal enhancer of mouse *Pou5f1/Oct4* (Hotta et al. 2009) (**Fig. S1G**). Following puromycin selection and clonal analysis via genomic PCR, a clone named NaiveRep was selected, which contained both *DPPA2^mKO2^* and *EOS^TagBFP^*. NaiveRep exists in two forms: NaiveRep_KF cells and NaiveRep_VAL, the latter obtained after culturing NaiveRep_KF cells in VALGöX medium for 48 h (**Fig. 1F**). Interestingly, NaiveRep_VAL cells expressed the fluorescent reporter EOS-TagBFP at a slightly higher level than NaiveRep_KF cells. In contrast, no difference in mKO2 fluorescence level was observed between the two cell types (**Fig. 1G**). Moreover, NaiveRep_VAL cells cultivated in VALGöX for 14 days exhibited morphological differentiation similar to B19_VAL cells. Overall, these findings confirm the VALGöX culture medium’s capacity to initiate conversion to naïve pluripotency, but also highlight its inability to complete the process.

### cDNA library screening

A library of 25 human cDNAs was generated using the W10 simian immunodeficiency virus (SIV)-based backbone lentiviral vector (Mangeot et al. 2000). The selected cDNAs encode transcriptional factors (*NANOG*, *SOX2*, *KLF2*, *KLF4*, *KLF5, KLF12, ESRRB*, *ESRRG*, *GBX2*, *TFCP2L1*, *NR5A2*, *SALL4*, *MYC*, and *GFI1*), a signaling molecule (ERAS), histone modification enzymes and chromatin modifiers (*PRMT6*, *KHDC1*, *KAT2B*, *KDM4D*, *SUV39H1*, BMI1, and *GASC1*), a cell-cycle protein (*CCNE1*), and viral oncogenes including SV40 Large T antigen (AgT) and adenovirus E1A-12S. Many of the selected genes have been demonstrated to support naïve pluripotency or facilitate the resetting of naïve-like features when overexpressed in mice and humans (Cartwright et al. 2005; Ema et al. 2008; Jiang et al. 2008; Parisi et al. 2008; Bourillot et al. 2009; Guo et al. 2009; Silva et al. 2009; Guo and Smith 2010; Han et al. 2010; Lee et al. 2012; Nagamatsu et al. 2012; Martello et al. 2013; Tai and Ying 2013; Uranishi et al. 2013; Stuart et al. 2014; Takashima et al. 2014; Qiu et al. 2015; Pastor et al. 2018; Festuccia et al. 2021; Lea et al. 2021). Our RNAseq analyses of rabbit embryos confirmed that all but one gene (*GFI1*) were expressed in the ICM/early epiblast of day 3 and 4 rabbit blastocysts (Bouchereau et al., 2022) (**Fig. S2A**).

The relative titers of the 25 viruses were estimated by serial dilution (**Fig. S2B**). Minor differences were observed between lentiviruses. It should be noted that the KLF2 lentivirus had a significantly lower titer than all the others. To test the ability of B19 cells to be infected by several lentiviruses simultaneously, we used four W10 lentiviral vectors expressing GFP, mKO2, tagBFP, and Katushka, respectively, which showed that up to 70% of the transduced cells could infected by the four vectors (**Figs. S2C, S2D**). A multiplicity of infection (MOI) of 50 viruses/cell was used for cDNA library transduction (*i.e.* approximately 2 viruses/cDNA/cell). A total of 2.4 x 10^5^ NaiveRep cells were transduced with the cDNA library and further cultured for seven days (**Fig. 2A**). Nearly 5,000 colonies were examined under epifluorescence, and 27 exhibiting increased mKO2- and/or tagBFP-associated fluorescence were picked and amplified for further analysis (later called NaiveRep_KF clones). Flow cytometry analysis of tagBFP and mKO2 fluorescence revealed significant variations in the percentage of positive cells, ranging from 1% to 80% (tagBFP) and from 1% to 7% (mKO2) (**Fig. 2B**). Upon analyzing the genomic DNA of the 27 clones by PCR, it was found that the average number of different provirus integrations was 9.3 +/-3.6 (**Fig. 2C**). The most frequently observed proviral DNAs were *ESRRG* (24 clones), *ERAS* (23 clones), *KLF2* (22 clones), *BMI1* (18 clones), *PRMT6* (17 clones), *NANOG* (16 clones), *ESRRB* (16 clones), and *AgT* (15 clones). Out of the 63 possible combinations of three different proviruses among the eight most frequent ones, seven were observed in more than 50% of the 27 clones analyzed: *KLF2*/*ERAS*/*ESRRB* (14/27); *KLF2*/*ERAS*/*BMI1* (14/27); *ERAS*/*ESRRG*/*PRMT6* (14/27), *ERAS*/*ESRRB*/*ESRRG* (13/27); *ERAS*/*ESRRG*/*BMI1* (14/27); *KLF2*/*ERAS*/*PRMT6* (15/27); and *KLF2*/*ERAS*/*ESRRG* (17/27). Interestingly, among the six clones (#8, #15, #17, #19, #22, and #26, respectively) with a percentage of cells exhibiting tagBFP expression greater than 30%, four clones harbored *KLF2*, *ERAS*, *ESRRG*, and *PRMT6* cDNAs ((#8, #15, #22, and #26, respectively), one clone harbored *KLF2*, *ERAS*, and *ESRRG* (#17), and one clone harbored *ERAS* and *ESRRG* (#19). In summary, the screen identified a combination of four provirus integrations (*KLF2*, *ERAS*, *ESRRG*, and *PRMT6*) with the highest frequency of occurrence among EOS-tagBFP-induced clones.

**Figure 2:**
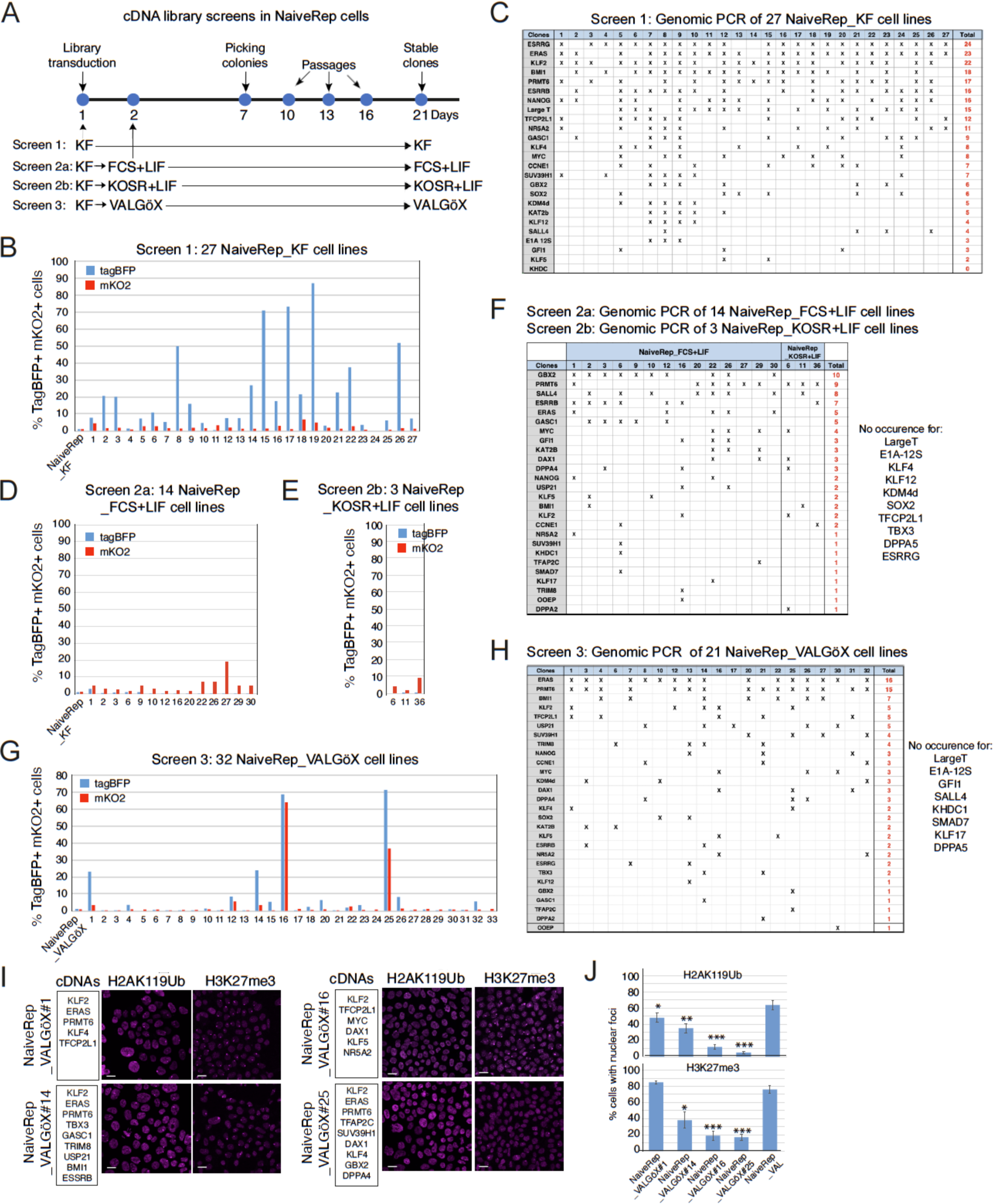
cDNA library screening. (**A**) Experimental scheme with four different screening conditions. (**B**) Flow cytometry analysis of EOS-tagBFP and DDPA2-mKO2 expression in 27 NaiveRep_KF cell lines. (**C**) Genomic PCR analysis showing proviral integrations in 27 NaiveRep_KF cell lines. (**D**) Flow cytometry analysis of EOS-tagBFP and DDPA2-mKO2 expression in 14 NaiveRep_FCS+LIF cell lines. (**E**) Flow cytometry analysis of EOS-tagBFP and DDPA2-mKO2 expression in three NaiveRep_KOSR+LIF cell lines. (**F**) Genomic PCR analysis showing proviral integrations in the 14 NaiveRep_FCS+LIF cell lines and three NaiveRep_KOSR+LIF cell lines. (**G**) Flow cytometry analysis of EOS-tagBFP and DDPA2-mKO2 expression in 32 NaiveRep_VALGöX cell lines. (**H**) Genomic PCR analysis showing the cDNAs found in 21 of the 32 NaiveRep_VALGöX cell lines. (**I**) Immunostaining of H2AK119Ub and H3K27me3 histone marks in four NaiveRep_VALGöX cell lines. Scale bars, 20 µm. (**J**) Histograms of the percentage of cells with H2AK119Ub and H3K27me3 nuclear foci in four NaiveRep_VALGöX cell lines and NaiveRep_KF control cells (T-test compared to control cells: *, p<0.05; **, p<0.01; ***, p<0.001).

In the second screening experiment, 2.4 x 10^5^ NaiveRep cells were transduced with an updated cDNA library containing 11 additional human cDNAs (*SMAD7*, *DPPA2*, *DPPA4*, *DPPA5*, *KLF17*, *TBX3*, *DAX1*, *TRIM8*, *USP21*, *TFAP2C*, and *OOEP*). The MOI was reduced to 15 viruses/cell (*i.e.* approximately 0.4 virus/cDNA/cell) to decrease the average number of cDNAs per clone, thus increasing the screening stringency. The transduced cell population was shifted from FGF2/KSR to either fetal calf serum (FCS)+LIF (screen 2a) or KOSR+LIF (screen 2b) (**Fig. 2A**). Note that in these two culture conditions, unmodified NaiveRep cells were unable to self-renew and differentiated. Approximately 5,000 colonies were examined under epifluorescence, with 14 colonies (screen 2a; **Fig. 2D**) and three colonies (screen 2b; **Fig. 2E**) exhibiting tagBFP- and/or mKO2-associated fluorescence. These colonies were picked and amplified. The average number of proviral integrations was 4.8 +/-2.4. The most frequently observed proviral integrations were *GBX2* (10/17 clones), *PRMT6* (9/17 clones), *SALL4* (8/17 clones), and *ESRRB* (7/10 clones) (**Fig. 2F**). No specific combination of proviral integrations could be identified in screens 2a and 2b.

In the third screening experiment, 2.4 x 10^5^ NaiveRep cells were transduced with the extended cDNA library using a MOI of 15. After 24 hours, the cells were transferred from FGF2/KSR to VALGöX culture medium and cultured for an additional 7 days (**Fig. 2A**). Almost 9,000 colonies were examined under epifluorescence, and 32 of them displayed stronger tagBFP- and mKO2-associated fluorescence. These colonies were picked and amplified for further analysis (later named NaiveRep_VALGöX clones). In these 32 cell clones, the percentage of fluorescent cells ranged from 1% to 71% (tagBFP) and from 1% to 64% (mKO2) (**Fig. 2G**). Of these 32 cell clones, 21 were analyzed for proviral integrations. The average number of integrations was 4.8 +/-2.0. The most frequently observed proviral integrations were *ERAS* (16 of 21 clones) and *PRMT6* (15 of 21 clones) (**Fig. 2H**). Notably, all clones exhibiting high EOS^tagBFP^ and DPPA2^mKO2^ expression contained the *KLF2* provirus (clones #1, #12, #14, #16, and #25) and all but one (clone #16) also harbored the *ERAS* and *PRMT6* proviruses. Consequently, the third screen allowed us to identify a common combination of three proviruses (*KLF2*, *ERAS* and *PRMT6*) in clones exhibiting higher expression of both EOS^tagBFP^ and DPPA2^mKO2^ naïve markers.

Two important insights can be drawn from the three screening experiments. First, the two fluorescent reporters were differentially activated across the screens. EOS^tagBFP^ was predominantly activated in the KOSR/FGF screening, while DPPA2^mKO2^ was preferentially activated in the KOSR/LIF and FCS/LIF screenings. Only in the VALGöX screening were both fluorescent reporters activated. This observation led us to utilize the VALGöX culture condition for subsequent experiments. Second, the most enriched proviruses differed from one screening to the other. *KLF2*, *ESRRG*, *ESRRB*, and *NANOG* were either poorly enriched or not enriched at all in the second and third screens. Conversely, *GBX2* and *SALL4* were exclusively enriched in the second screen, while *ERAS* was significantly more enriched in the first and third screens. Collectively, these findings indicate that the selection pressure on cDNAs is influenced by the culture regimen.

The four cell lines that displayed activation of both fluorescent reporters in the VALGöX culture condition (NaiveRep_VALGöX#1, NaiveRep_VALGöX#14, NaiveRep_VALGöX#16, and NaiveRep_VALGöX#25; **Figs 2G, S2E**) were chosen for further analysis. All four cell lines exhibited consistent growth for at least six passages (**Fig. S2F**). Immunofluorescence analysis of histone marks H3K27me3 and H2AK119Ub showed a reduction in the percentage of cells with nuclear foci, suggesting reactivation of the second X chromosome in some cells (H2AK119Ub: 48%, 35%, 11%, and 5% in #1, #14, #16, and #25, respectively, compared with 63% in control NaiveRep cells; H3K27me3: 38%, 19%, and 17% in #14, #16, and #25, respectively, compared with 76% in control NaiveRep cells; **Figs. 2I, 2J**). Notably, a positive correlation was observed between the percentage of EOS^tagBFP^-positive cells and the percentage of cells lacking nuclear foci H3K27me3 and H2AK119Ub (**Fig. S2G**), further supporting the naïve-like characteristics of the fluorescent cell population.

NaiveREP_KF#22 cells, obtained in the first screening using the KOSR+FGF culture regimen, were switched to VALGöX culture conditions for 7 days, generating NaiveRep_KF_VALGöX#22 cells. They contained six proviral integrations, *KLF2*, *ERAS*, *PRMT6*, *ESRRB*, *NANOG*, and *TFCP2L1*. The NaiveRep_KF#22 and NaiveRep_ KF_VALGöX#22 pair was used to investigate the relative contribution of the proviral integrations and VALGöX medium to chromatin remodeling. A significant decrease in the repressive mark H3K9me3 and an increase in the permissive mark H3K14Ac were observed in NaiveRep_KF_VALGöX#22 cells when compared to NaiveRep and NaiveRep_KF#22 cells (**Figs. S2H, S2I**). Additionally, the percentage of cells lacking H2AK119Ub nuclear foci further decreased in NaiveRep_KF_VALGöX#22 cells compared to NaiveRep_KF#22 cells. Overall, these observations suggest that both the proviruses and VALGöX culture medium are required for chromatin remodeling.

### Stabilization of a naïve-like pluripotent state using KLF2/ERAS/PRMT6 gene cocktail and VALGöX culture regimen

To investigate the role of KLF2 (K), ERAS (E), and PRMT6 (P) individually or in combinations in restoring naïve pluripotency characteristics in rabbit iPS cells, B19 iPSCs were transfected with one, two, or three of the plasmids expressing V5-tagged KLF2 and neoR (*pKLF2:V5-neo*), HA-tagged ERAS and hygroR (*pHA:ERAS-hygro*), and Flag-tagged PRMT6 and puroR (*pFlag:PRMT6-puro*), respectively. Following single, double, and triple antibiotic selection in VALGöX medium, resistant colonies were pooled and further propagated for 10 passages before phenotypic analysis (**Fig. S3A**). Similar to control cells (B19_VAL, **Fig. 1C**), B19_VAL cells expressing *ERAS* and *PRMT6* transgenes individually or in combination (referred to as E, P, and EP, respectively), gradually differentiated between passage 5 (P5) and P10 (**Fig. 3A, S3B**). In contrast, B19_VAL cells expressing the *KLF2* transgene, either alone or with *ERAS* and *PRMT6* transgenes (K, KE, KP, and KEP, respectively), successfully overcame the P5 to P10 transition and continued self-renewing without visible differentiation. This indicates that KLF2 is crucial for the long-term self-renewal of B19 cells cultivated in VALGöX. Interestingly, cells expressing the three transgenes (KEP) exhibited higher expression of the naïve pluripotency cell surface marker CD75 (Collier et al. 2017; Trusler et al. 2018) compared to K, KE, KP (**Fig. 3A, 3B**). The KEP cell population also had fewer cells with nuclear foci after immunostaining for H3K27me3 and H2AK119Ub compared to K, KE, KP, suggesting higher rate of X chromosome reactivation when *KLF2*, *ERAS*, and *PRMT6* were co-expressed (**Fig. 3A, 3C**). Furthermore, double immunostaining of CD75/H3K27me3 revealed a strong correlation between CD75 expression and the reactivation of the second X chromosome (**Fig. 3D**).

**Figure 3:**
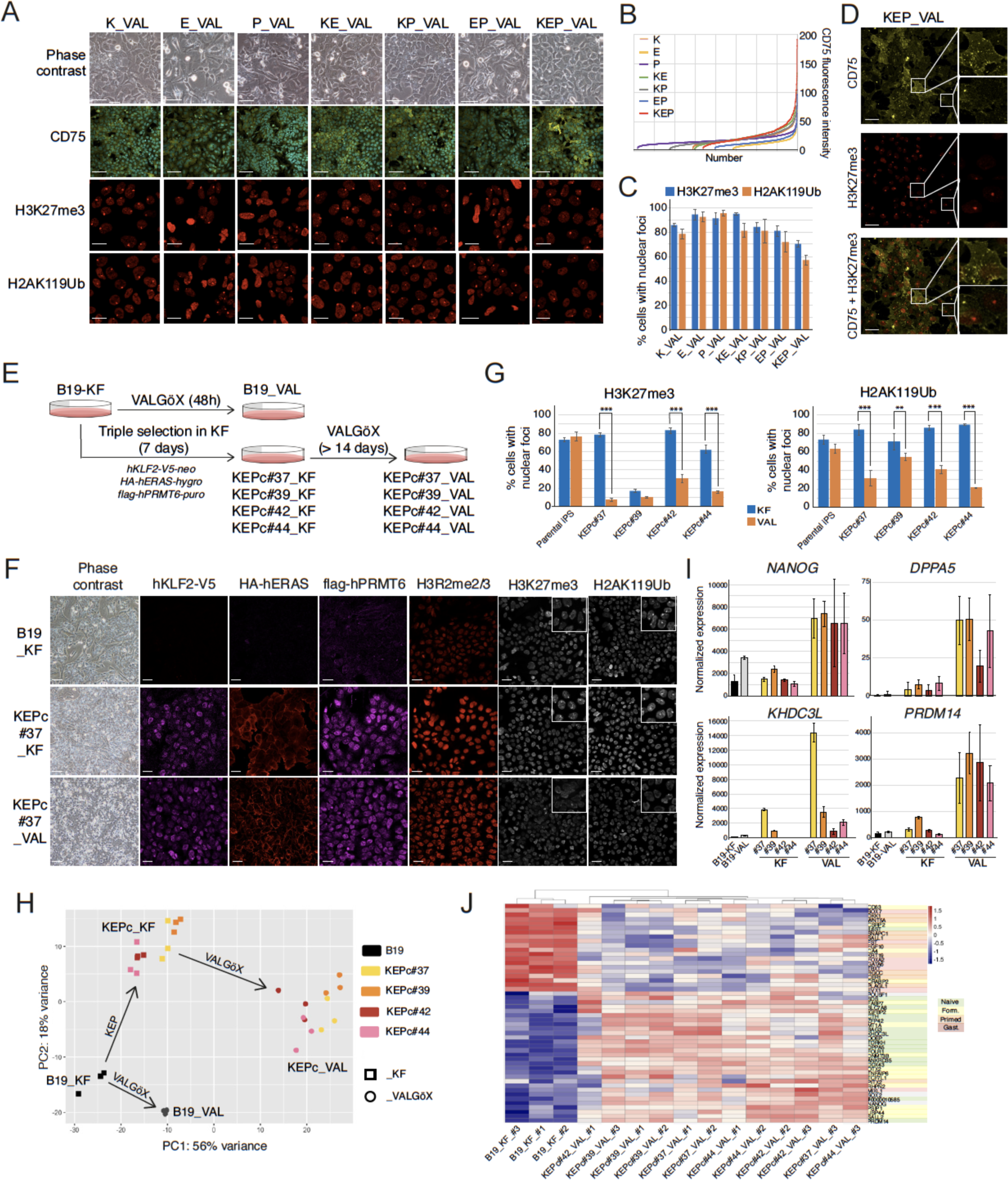
Stabilization of a naïve-like pluripotent state with KLF2/ERAS/PRMT6 gene cocktail. (**A**) Phase contrast and confocal images of B19_VAL before and after single, double, and triple transfection with PiggyBac plasmids expressing KLF2-V5 (K), HA-ERAS (E), and flag-PRMT6 (P). Immunostainings with CD75, H3K27me3 and H2AK119Ub antibodies are displayed. Scale bars: phase contrast, 100µm; CD75, 50µm; histone modifications, 20µm. (**B**) Graphical representation of the number of cells displaying fluorescence intensities between 0 and 200 (arbitrary units) after immunostaining of CD75. (**C**) Percentage of cells exhibiting H3K27me3 and H2AK119Ub nuclear foci in control, single-, double-, and triple-transfected cell populations. (**D**) Immunostaining of KEP_VAL cells with CD75 and H3K27me3 antibodies. Scale bar: 40µm. (**E**) Experimental design for generating KEPc_KF and KEPc_VAL cell lines. (**F**) Phase contrast and confocal images of KLF2-V5, HA-ERAS, flag-PRMT6, H3R2me2/3, H3K27me3, and H2AK119Ub in control B19_KF, KEPc#37_KF, and KEPc#37_VAL cells. Scale bars: phase contrast, 100µm; immunostainings, 30µm. (**G**) Percentage of cells containing H3K27me3 and H2AK119Ub nuclear foci in control parental B19 iPSCs, KEPc_KF, and KEPc_VAL cell lines (T-test comparison of KF and VAL conditions: **, p<0.01; ***, p<0.001). (**H**) Two-dimensional PCA of the transcriptomic data obtained from five cell populations (B19, KEPc#37, KEPc#39, KEPc#42, KEPc#44) into two media (Square, KF; Circle, VALGöX) as indicated (three biological replicates). (**I**) Histograms illustrating gene expression in B19_KF, B19_VAL_48h, KEPc_KF, and KEPc_VAL cell lines (based on RNA-seq data). (**J**) Heatmap representation of differentially expressed genes between B19_KF, KEPc#37_VAL, KEPc#39_VAL, KEPc#42_VAL, and KEPc#44_VAL cells (genes are selected from the list of 80 differentially expressed genes between inner cell mass (E3.5), early epiblast (E4.0), mid-epiblast (E5.0) and late epiblast (E6.0 and E6.6) of rabbit embryos (Bouchereau et al. 2022).

To further investigate the synergistic effect of KEP transgenes and the VALGöX culture regimen in reprogramming iPS cells, B19 cells cultivated in KF were transfected with *pKLF2:V5-neo*, *pHA:ERAS-hygro* and *pFlag:PRMT6-puro*. Four triple-resistant clones, hereafter referred to as KEPconstitutive (KEPc)#37_KF, KEPc#39_KF, KEPc#42_KF, KEPc#44_KF, were isolated. These clones were then switched to VALGöX culture conditions and further cultivated for 20 days, resulting in KEPc#37_VAL, KEPc#39_VAL, KEPc#42_VAL, and KEPc#44_VAL cells, respectively (**Fig. 3E**). Immunofluorescence analysis revealed membrane-bound expression of HA:ERAS and nuclear expression of KLF2:V5 and Flag:PRMT6 in both KEPc_KF and KEPc_VAL cells (**Figs. 3F, S3C**). The H3R2me3 mark levels were significantly increased, reflecting the arginine methyltransferase protein activity of PRMT6. AKT phosphorylation was also markedly increased, reflecting ERAS activity (**Fig. S3D)**(Takahashi et al. 2003), whereas ESRRB expression, a target gene KLF2, was higher in KEPc cells compared to control cells (**Fig. S3F**). The KF to VAL conversion substantially reduced the ratio of cells with nuclear foci observed after immunostaining for histone marks H3K27me3 and H2AK119Ub, suggesting the reactivation of the second X chromosome in the majority of KEPc_VAL cells (**Figs. 3G, S3C**). Overall, these findings demonstrate that the expression of KLF2, ERAS, and PRMT6, in conjunction with the VALGöX culture regimen, promotes epigenetic remodeling consistent with naïve-like pluripotency.

To further validate the transition from primed to naïve-like pluripotency states, we analyzed the transcriptomes of B19_KF, B19_VAL, KEPc_KF, and KEPc_VAL cell lines using RNA sequencing. Upon analysis of these PSC lines, four distinct clusters emerged in a principal component analysis (PCA). The least significant difference was observed between B19_KF and B19_VAL, with 597 differentially-expressed genes (DEGs), while the most significant difference was found between B19_KF cells and KEFc_VAL cells, with 1,707 DEGs (**Figs. 3H**, **S3E**). In KEPc_VAL cells, the expression of naïve pluripotency genes *DPPA2*, *DPPA5*, *PRDM14*, *KHDC3L*, *FOLR1*, *ST6GAL1 (CD75)*, *TDRKH*, and *DNMT3B* increased compared to B19_KF and B19_VAL, while the expression of primed pluripotency and gastrulation genes *CER1*, *DKK1*, *TBXT*, and *EVX1* decreased (**Figs. 3I**, **S3F**). Interestingly, some naïve pluripotency, such as *DPPA5*, *PRDM14*, and *KHDC3L*, exhibited a strong increase in expression in KEPc_VAL cells compared to all other cell types, including KEPc_KF. This further supports the synergistic effect of KEP transgenes and the VALGöX culture regimen.

Finally, a heatmap of differentially expressed genes between B19_KF and KEFc_VAL samples was calculated based on a list of 80 differentially expressed genes between inner cell mass (E3.5), early epiblast (E4.0), mid-epiblast (E5.0) and late epiblast (E6.0 and E6.6) of rabbit embryos (Bouchereau et al. 2022) (**Fig. 3J**). Fifty of these genes were found to be differentially expressed between B19_KF and KEFc_VAL cells. Notably, most of the genes overexpressed in B19_KF cells were predominantly expressed at E6 and E6.6 (referred to as “primed” and “gastrulation”), whereas the vast majority of the genes overexpressed in KEPc_VAL cells were predominantly expressed at E3.5 and E4.0 (referred to as “formative” and “naïve”). Overall, these results indicate that the expression of *KLF2*, *ERAS*, and *PRMT6* in iPSCs cultured in VALGöX facilitates their conversion from primed to formative/naive-like pluripotency.

### Enhanced colonization of rabbit embryos by iPSCs overexpressing KLF2, ERAS and PRMT6 in VALGöX culture condition

We next investigated the ability of KEPc to colonize rabbit embryos. KEPc#37_KF, KEPc#37_VAL, KEPc#44_KF, and KEPc#44_VAL cells were injected into early morula stage embryos at a ratio of eight cells per embryo. The injected embryos were then cultured for three days and developed into late blastocysts (3 days in vitro, DIV) (**Fig. 4A, S4**). In three independent experiments, 63% of the blastocysts displayed GFP^+^ cells following the injection of KEPc_VAL cells, compared to 23% after injecting KEPc_KF cells and only 5% after injecting parental iPS B19_KF cells (**Fig. 4B**). A significant difference was observed in the number of GFP^+^ cells among the three cell types. The majority of embryos injected with eight parental B19_KF or KEPc_KF cells had fewer than five GFP^+^ cells after 3 DIV. Only three embryos (7%) injected with KEPc_KF cells had more than five (but fewer than 10) cells. In contrast, embryos injected with eight KEPc_VAL cells exhibited notably increased colonization. For KEPc#37_VAL, among GFP+ embryos, up to 66% of them contained over 10 GFP^+^ cells, and 50% had more than 20 GFP^+^ cells. The vast majority of the GFP^+^ cells expressed SOX2 (**Fig. 4A, S4**). Overall, these results indicate that the KEP transgenes, when combined with the VALGöX culture regimen, significantly enhance both the growth and embryo colonization capabilities of iPS cells.

**Figure 4:**
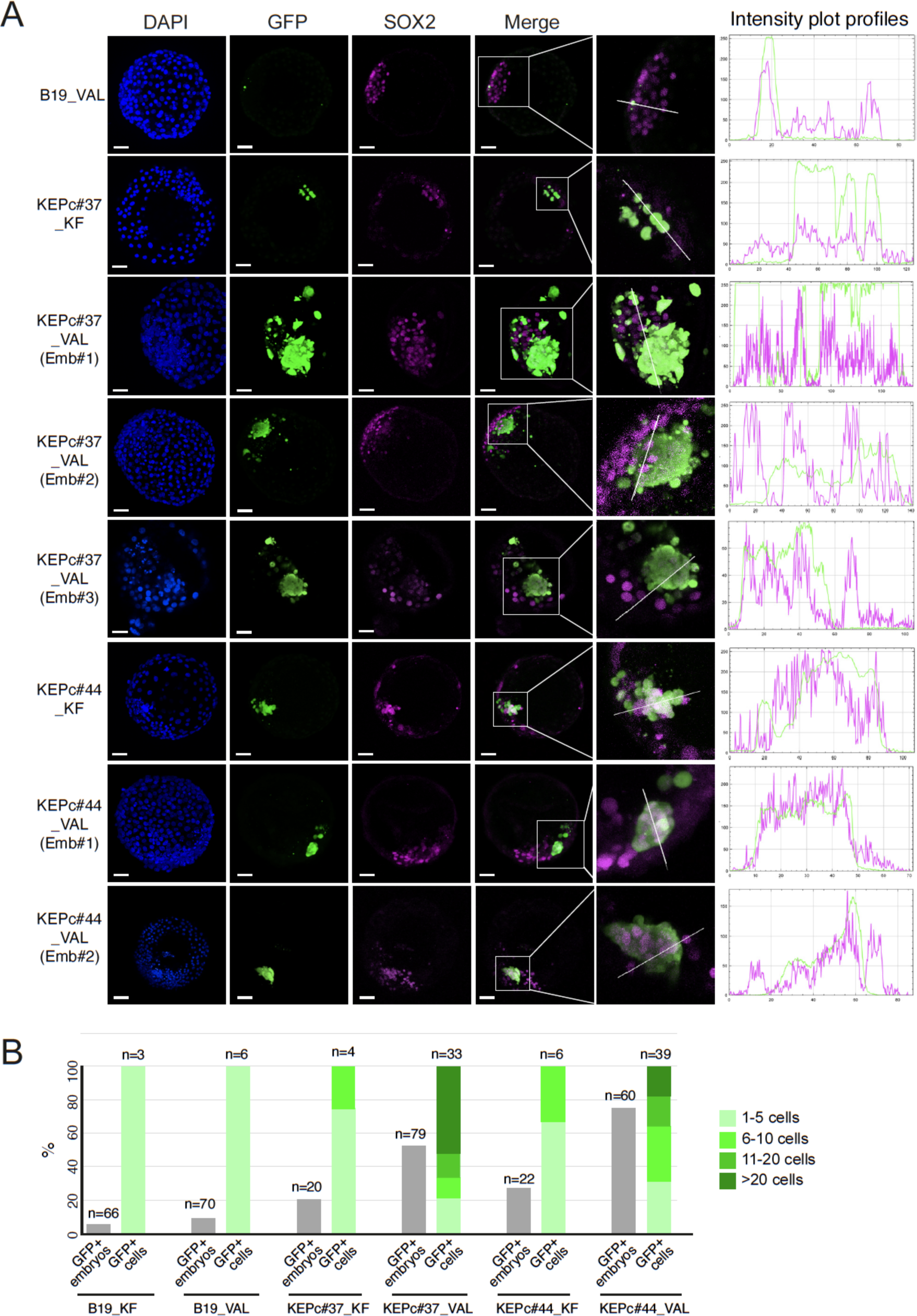
Enhanced colonization of rabbit embryos by iPSCs overexpressing KLF2, ERAS and PRMT6. (**A**) Confocal images of late-blastocyst-stage rabbit embryos (E5.0, 3DIV) acquired after the microinjection of B19_KF, KEPc#37_KF, KEPc#37_VAL, KEPc#44_KF, and KEPc#44_VAL cells into early morula-stage (E2.8) embryos (scale bars: 50 µM). On the right part of the panel, a single line has been drawn through the ICM of the embryos for further intensity profile measurement using Fiji. The resulting fluorescence intensity plot profile across multiple channels (green for GFP signal, purple for SOX2 signal) demonstrates that GFP cells are also SOX2^+^. (**B**) Percentage of rabbit embryos with GFP^+^ cells and the distribution according to GFP^+^ cell number among GFP^+^ embryo.

### Transcriptomic signature of iPSCs associated with enhanced embryonic colonization

Our goal was to compare the transcriptomes of B19_KF, B19_VAL, KEPc_KF and KEPc_VAL cells in order to identify potential molecular determinants of embryonic colonization capacity. Given that KEPc_VAL cells displayed a significantly higher embryonic colonization capacity compared to B19_KF, B19_VAL, and KEPc_KF cells, we aimed to pinpoint genes that were up-or down-regulated due to the synergy between KEP transgenes and VALGöX culture conditions, focusing our analysis on these genes. We excluded genes whose expression was altered by either factor alone from our analysis. To achieve this, we removed the DEGs identified between B19_KF and B19_VAL (representing the “VALGöX effect”) and those between B19_KF and KEPc_KF (representing the “KEP effect”) from the list of 1707 DEGs identified between B19_KF and KEPc_VAL (**Fig. 5A**). We found that “VALGöX effect” and “KEP effect” DEGs accounted for 23% and 0.4% of all DEGs, respectively. After this subtraction, we were left with 1,317 genes that represented the combined “KEP+VALGöX synergetic effect”, with 783 being up-regulated and 534 down-regulated (**Fig. 5B**). We next investigated the expression profile of these “KEP+VALGöX synergetic effect” genes in rabbit preimplantation embryos to determine if they were enriched at specific developmental stages. To this end, we extracted the 100 most up-regulated genes and the 100 most down-regulated genes from the list of 1,317 DEGs. Utilizing our unique transcriptome dataset (Bouchereau et al. 2022), we analyzed their expression in rabbit preimplantation embryos and visualized the results in a heatmap (**Fig. 5C**). Our analysis revealed that many down-regulated genes, such as *DKK3*, *HDACD2*, and *MPP6* exhibited higher expression in the epiblast of E6.0/E6.6 embryos. Conversely, up-regulated genes such as *DPPA3*, *DPPA5*, *KHDC3L*, *SOX15,* and *ZFP42* displayed higher expression in morula and epiblast of E3.5-E5.0 embryos. These findings suggest that the synergy between KEP transgenes and the VALGöX culture regimen led to the up-regulation of genes active in the early epiblast.

**Figure 5:**
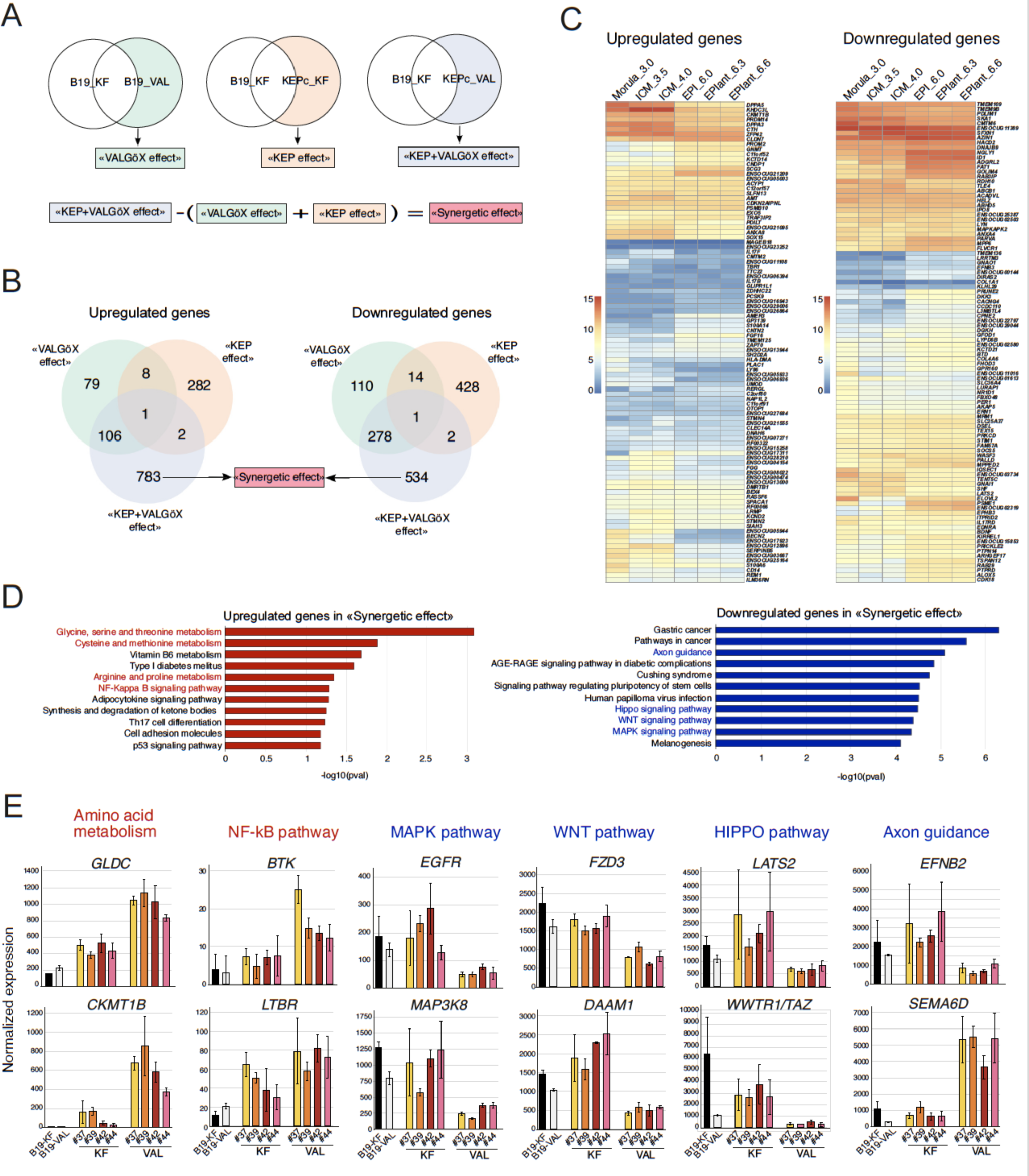
Transcriptomic signature of iPSCs associated with enhanced embryonic colonization. (**A**) Logic diagram for delineating the genes activated or repressed by the synergistic effect of VALGöX culture medium and KEP transgenes. (**B**) Venn diagrams displaying the number of up- and down-regulated genes in three gene lists: “VALGöX effect”, “KEP effect”, and “KEP+VALGöX synergetic effect”. (**C**) Heatmap representation of the expression levels in rabbit preimplantation embryos for the 200 most differentially expressed genes (100 most up- and 100 most down) in the “KEP+VALGöX synergetic effect” list (E, embryonic day; EPI, epiblast; EPIant, anterior epiblast; EPIint, intermediate epiblast; EPIpost, posterior epiblast). To facilitate visual presentation, ENSOCUG numbers were shortened by removing 000000 as follows: ENSOCUGxxxxx (**D**) KEGG pathway enrichment analysis based on the list of up- and down-regulated genes associated with the “KEP+VALGöX synergetic effect”. (**E**) Histogram representation of the normalized expression levels of selected genes in B19_KF, B19_VAL_48h, KEPc_KF, and KEPc_VAL cells (from RNA-seq data).

The KEGG pathway enrichment analysis, conducted using the list of 783 upregulated genes associated with the “KEP+VALGöX synergetic effect”, revealed connections to molecular pathways involved in amino acid metabolism (**Figs. 5D, 5E, S5**). Specifically, this included the activation of *SDS* (serine dehydratase), *CTH* (cystathionine γ-lyase), *GLDC* (glycine decarboxylase), *PSAT1* (phosphoserine aminotransferase), *AMT* (aminomethyltransferase), *ALDH7A1/8A1* (aldehyde dehydrogenase 7A1 and 8A1), *GNMT* (glycine N-methyltransferase), *GSS* (glutathione synthetase), *SMS* (spermine synthase), *CNDP1* (carnosine dipeptidase 1), *CKMT1B* (creatine kinase mitochondrial 1B), *ACYP1* (Acylphosphatase 1), and *PYCR2* (pyrroline-5-carboxylate reductase 2). These enzymes regulate serine, glycine, lysine, cysteine, alanine, arginine, methionine, proline, tryptophane, and glutamate homeostasis, as well as processes like pyruvate production (*SDS*), one-carbon metabolism (*SDS*, *GLDC*, *PSAT1*, and *GNMT*) and energy metabolism (*CKMT1B*). In addition, the analysis highlighted the involvement of the NF-κB signaling pathway, specifically through the activation of *BTK* (Bruton’s tyrosine kinase), *EDARADD* (Ectodysplasin A receptor), *ZAP70* (zeta-chain of T cell receptor associated protein kinase 70), *LTBR* (lymphotoxin-β receptor), and *CD14* (coreceptor for lipopolysaccharides). These five factors can activate IKK degradation and subsequent NF-kB pathway activation. Furthermore, the KEGG pathway enrichment analysis revealed a strong association with the p53 signaling pathway. This association includes the activation of *p53R2*/*RRM2B* (ribonucleotide reductase regulatory TP53 inducible subunit 2B) and *GADD45G* (Growth Arrest and DNA Damage-inducible 45γ), which encode two p53 effectors involved in DNA repair and damage prevention. Additionally, the analysis identified the activation of *SESN2* (sestrin 2), an inducer of MDM2 degradation and subsequent p53 activator. Overall, these results suggest that KEPc_VAL cells underwent significant changes in amino acid and one-carbon metabolism, as well as in enhancing cell survival, and protecting against DNA damage.

The KEGG pathway enrichment analysis conducted using the list of 534 downregulated genes associated with the “KEP+VALGöX synergetic effect” revealed strong connections with MAPK, WNT, and Hippo signaling pathways (**Figs. 5D, 5E, S5**). With regard to the MAPK signaling pathway, it involved the downregulation of several molecules from the ERK, JNK, and p38 MAPK pathways, including receptor-encoding genes *PDGFRB*, *EGFR*, and *TGFBR2*; signaling molecule-encoding genes *ARAF* (RAF-A), *MAP3K8*, and *MAPKAPK2*; MAPK phosphatase-encoding gene *DUSP2*; and the ERK/p38-target gene *ELK-4*. In terms of the WNT signaling pathways, it involved the downregulation of several genes encoding WNT ligands and modulators (*WNT5B* and *GPC4*), their receptor (*FZD3*/*4*), and adaptors (*PRICKLE2*, *AXIN2*, and *DAAM1*). *PPP3CA* (calcineurin A) was also downregulated, involved in the non-canonical WNT/Ca2^+^ signaling pathway. Regarding the HIPPO signaling pathway, *WWC1*/*KIBRA*, *LATS2*, and *PPP2R2B*, which code for upstream regulators of YAP/TAZ activity, were downregulated. Finally, “Axon guidance” was strongly enriched in the KEGG pathway analysis of downregulated genes. This involved the downregulation of *EFNB2* (Ephrin-B2), *EPHB1*, *EPHB3*, and *EPHA4* (EPH-related receptors). *NTN1* (Netrin-1) and *PLXNA2* (Plexin A2) were also downregulated. Collectively, these results suggest that KEPc_VAL cells attenuate the activity of ERK, JNK, p38MAPK, YAP/TAZ, Ephrin, as well as both canonical and non-canonical WNT signaling pathways.

### KLF2, ERAS and PRMT6 hold iPSCs in a naïve-like, colonization-competent state

We sought to determine if the enhanced colonization ability of KEPc_VAL cells in host embryos resulted from stable reprogramming to the naïve state, or was merely a consequence of KEP transgene expression. To investigate this, we designed iPSCs cells to express *KLF2*, *ERAS* and *PRMT6* transgenes in an inducible manner. We transfected B19_KF cells with three plasmids encoding Shield1-activatable DD:KLF2:V5 (*pDD:KLF2:V5-neo*), DD:HA:ERAS (*pDD:HA:ERAS-hygro*), and DD:Flag:PRMT6 (*pDD:Flag:PRMT6-puro*), respectively. These vectors contained mutated FKBP12-derived destabilization domain (DD) fused to KLF2:V5, HA:ERAS, and flag:PRMT6 coding sequences, enabling interaction and stabilization by Shield1 (Banaszynski et al. 2006). We transfected B19_KF cells without Shield1 and identified two triple-resistant cell lines, KEPi#13_KF and KEPi#18_KF. We then transferred these cell lines to VALGöX medium supplemented with Shield1 and cultured them for three weeks, producing KEPi#13_VAL+S and KEPi#18_VAL+S (**Fig. 6A**). In another experiment, we transfected B19_KF cells with the three plasmids expressing Shield1-activatable KLF2, ERAS, and PRMT6 and immediately transferred them to VALGöX medium with 1000 nM Shield1 before culturing for at least three weeks, yielding KEPi#28_VAL+S and KEPi#36_VAL+S cells. We then cultured KEPi#13_VAL+S, KEPi#18_VAL+S, KEPi#28_VAL+S, and KEPi#36_VAL+S in Shield1-free medium for seven days, resulting in KEPi#13_VAL+S-S, KEPi#18_VAL+S-S, KEPi#28_VAL+S-S, and KEPi#36_VAL+S-S. There was almost complete silencing of ERAS and PRMT6 and partial silencing of KLF2 after Shield1 withdrawal (**Figs. 6B, S6A**). The H3R2me2/3 histone mark and ph-AKT levels decreased (**Figs. S6A**, **S6C, S6D**). However, despite Shield1 being withdrawn, the number of cells with nuclei foci did not change, suggesting that the XaXa status of KEPi cells was not altered (**Fig. S6B**). We analyzed the transcriptomes of KEPi#_KF, KEPi#_VAL+S, and KEPi#_VAL+S-S cell lines and visualized the data using two-dimensional PCAs (**Fig. 6C**). Transgene activation in VALGöX culture conditions caused significant transcriptome changes in the four analyzed clones. Transcriptome changes induced by Shield1 withdrawal were minor in KEPi#13 cells, moderate in KEPi1#18 cells, and substantial in KEP#28 and KEPi#36 cells. Naïve pluripotency markers *DPPA2*, *DPPA5*, *KHDC3L*, *KLF4*, *PRDM14*, and *ZFP42* were down-regulated, while primed pluripotency markers *SALL1*, *CDH2*, *FGF5*, *PLAGL1*, and *DKK1* were upregulated in KEPi#_VAL+S-S cells compared to KEPi#_VAL+S (**Figs. 6D, S6E**). This observation suggests that transgene silencing caused KEPi_VAL cells to depart from naïve-like pluripotency. Nevertheless, the Shield1-deprived cells continued to self-renew without any apparent differentiation.

**Figure 6:**
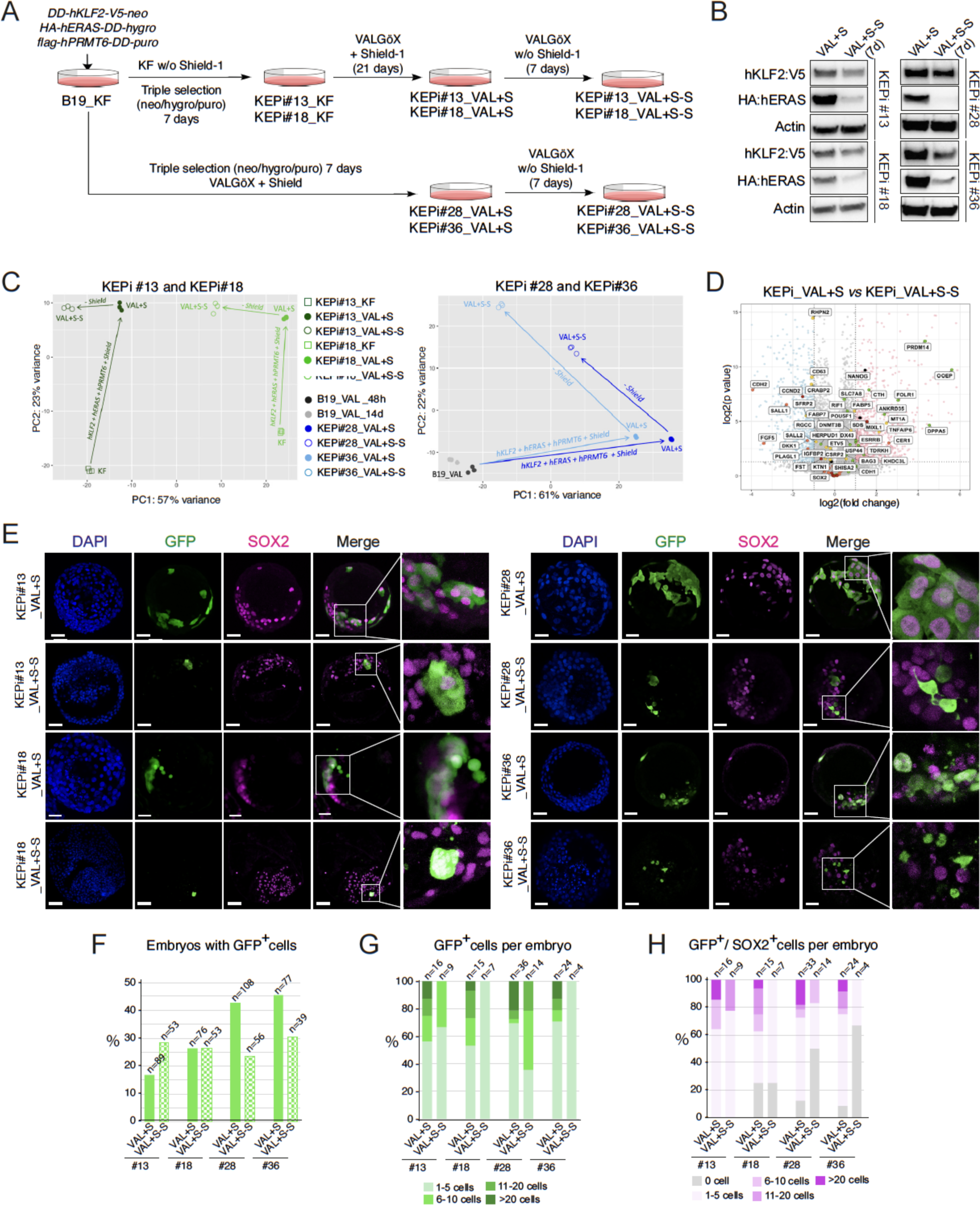
KLF2, ERAS and PRMT6 maintain iPSCs in a naïve-like, colonization-competent state. (**A**) Experimental scheme of KEPi_VAL+S and KEPi_VAL+S-S cell line generation. (**B**) Expression of KLF2-V5 and HA-ERAS analyzed through Western blotting in KEPi#13, KEPi#18, KEPi#28, and KEPi#36 cells before and after withdrawal of Shield1. (**C**) Two-dimensional PCA of the transcriptome of the indicated cell lines (three biological replicates). (**D**) Volcano plot representation of DEGs between KEPi_VAL+S and KEPi_VAL+S-S cells. (**E**) Confocal images of late-blastocyst-stage rabbit embryos (E5.0, 3DIV) acquired after microinjection of KEPi_VAL+S and KEPi_VAL+S-S cells into morula-stage (E2) embryos (scale bars: 50 µM). (**F**) Percentage of embryos with GFP^+^ cells with each cell line. (**G**) Distribution of embryos according to GFP^+^ cell numbers for each cell line. (**H**) Distribution of embryos according to GFP^+^/SOX2^+^ cell numbers for each cell line.

To assess the embryonic colonization capacity of KEPi_VAL+S and KEPi_VAL+S-S cells, we injected eight cells from each cell line into rabbit morulae. Embryos injected with KEPi_VAL+S cells were cultured in the presence of Shield1, while embryos injected with KEPi_VAL+S-S cells were cultured without inducer. Results were obtained from at least three independent experiments for each of the eight cell lines tested. A total of 35% of the analyzed blastocysts (n = 350) exhibited GFP^+^ cells after KEPi_VAL+S cell injection (25.8%, 26.3%, 42.6%, and 45.5% in #13, #18, #28, and #36 clones, respectively) compared to 25% (n = 201) after KEPi_VAL+S-S injection (28.3%, 26.4%, 19.6%, and 28.2% in #13, #18, #28, and #36 clones, respectively) (**Fig. 6E, 6F**). The GFP^+^ cells appeared organized in clusters or scattered throughout the embryo including the trophoblast. The average number of GFP^+^ cells per embryo was 6.5 (ranging from 1 to 27) after the injection of KEPi_VAL+S cells (**Figs. 6G**). The vast majority of GFP^+^ cells expressed SOX2 (5.8 cells/embryo, ranging from 1 to 27) and were localized in the epiblast (**Fig. 6H**). Rare SOX2^+^ cells were found in the TE (**Fig. 6E**). The average number of GFP^+^ and GFP^+^/SOX2^+^ cells decreased in embryos injected with KEPi_VAL+S-S cells (4.7 GFP^+^ cells per embryo, ranging from 1 to 10; 4.2 GFP^+^/SOX2^+^ per embryo, ranging from 1 to 10) (**Fig. 6G, 6H, S5**). Nevertheless, it remained substantially higher than the average number of GFP^+^ and GFP^+^/SOX2^+^ cells observed in embryos injected with parental B19_VAL cells (1.25 GFP^+^ per embryo, ranging from 1 to 2; one GFP^+^/SOX2^+^ per embryo, ranging from 0 to 2). Altogether, these results suggest that continuous transgene expression is required for preserving full embryonic colonization capability. Upon down-regulation of KLF2, ERAS and PRMT6, a substantial fraction of the cell population departs from naïve-like pluripotency and their embryonic colonization ability is subsequently reduced. We leveraged KEPi_VAL+S and KEPi_VAL+S-S cells to further investigate the transcriptomic characteristics of cells with high colonization potential. For this purpose, we identified the DEGs between the four KEPi_VAL+S cells on one side and the four KEPi_VAL+S-S cells on the other side (**Fig. S6F**). A KEGG pathway enrichment analysis of the 690 downregulated genes in the KEPi_VAL+S-S cells highlighted “Glycine, serine, and threonine metabolism” (**Fig. S6G**). This involved the downregulation of *ALDH7A1*, *CKMT1B*, and *CTH* genes (**Fig. S6H**), which were previously found upregulated in KEPc_VAL cells compared to B19_VAL cells (**Fig. 5E, S5**). In contrast, a KEGG pathway enrichment analysis of the 551 upregulated genes in KEPi_VAL+S-S cells highlighted WNT, MAPK, and HIPPO signaling pathways (**Fig. S6G**). This involved the upregulation of *EGFR*, *MAP3K8*, *PRICKLE2*, *LATS2*, and *WWTR1/TAZ* (**Fig. S6H**), which were previously found downregulated in KEPc_VAL cells compared to B19_VAL cells. Regarding the WNT signaling pathway, additional upregulated genes in KEPi_VAL+S-S cells included those encoding FZD7 and ROR1 receptors. Finally, “Axon guidance” was strongly enriched in the KEGG pathway analysis of KEPi_VAL+S-S cells. It involved the upregulation of genes encoding Ephrin B2 and Ephrin receptor B3, and the downregulation of gene encoding Semaphorin 6A. It can be concluded that the transcriptomic changes associated with the transition from KEPi_VAL+S to KEPi_VAL+S-S are a mirror image, albeit attenuated, of those associated with the transition from B19_VAL to KEPc. Interestingly, the four KEPi_VAL cell lines differed in this respect: the average +S/+S-S fold-change calculated from the 42 selected genes in Fig. S6G varied significantly between cell lines, with the minimum variation in KEPi#13_VAL and KEPi#18_VAL and the maximum in KEPi#28_VAL and KEPi#36_VAL cells (**Fig. S6I**). These differences in transcriptomic stability are consistent with the results of embryonic colonization, where the KEPi#28_VAL+S-S and KEPi#36_VAL+S-S cells displayed a reduced capacity for embryo colonization compared to the two other cell lines. Altogether, these results indicate that both lower expression of key regulators of MAPK, WNT, HIPPO, and Ephrin signaling pathways, and higher expression of key regulators of amino acid metabolism and NF-kB signaling pathway are associated with the enhanced ability of KEPc_VAL and KEPi_VAL+S cells to colonize the host epiblast.

### KEP_VAL cells with the highest CD75 expression exhibit massive colonization capacity

We had previously noted that a sub-population of B19_VAL_KEP cells expressed a high level of CD75 (**Fig. 3A**). Given this observation, we investigated whether a CD75^high^ cell sub-population exists in the KEPi_VAL+S cells and if these cells possess unique embryo colonization capabilities. To explore this, CD75^high^ cells were FACS-sorted from both KEPi#13_VAL+S and KEPi#28_VAL+S cell populations. They were then cultured for two distinct periods: 24 hours (referred to as condition I) and five days (referred to as condition II) before being injected into rabbit morulae (**Fig. 7A, 7B**). For embryos in condition I, 77% and 100% displayed GFP^+^ cells post-injection of KEPi#13_CD75^high^ and KEPi#28_CD75^high^ cells, respectively (**Fig. 7C, 7D**). Remarkably, 70% (KEPi#13) and 100% (KEPi#28) of these embryos had incorporated more than 20 GFP^+^ cells (**Figs**. **7E****, S7A**). In the case of condition II, 51% of the embryos showed GFP^+^ cells after being injected with KEPi#13_CD75^high^ and 100% after injection with KEPi#28_CD75^high^ cells. Additionally, 32% of the embryos from the KEPi#13 group and 63% from the KEPi#28 group incorporated over 20 GFP^+^ cells. These findings indicate that the CD75^high^ cells present in the KEPi_VAL+S cell populations exhibit a considerably enhanced ability for both epiblast and trophoblast colonization.

**Figure 7:**
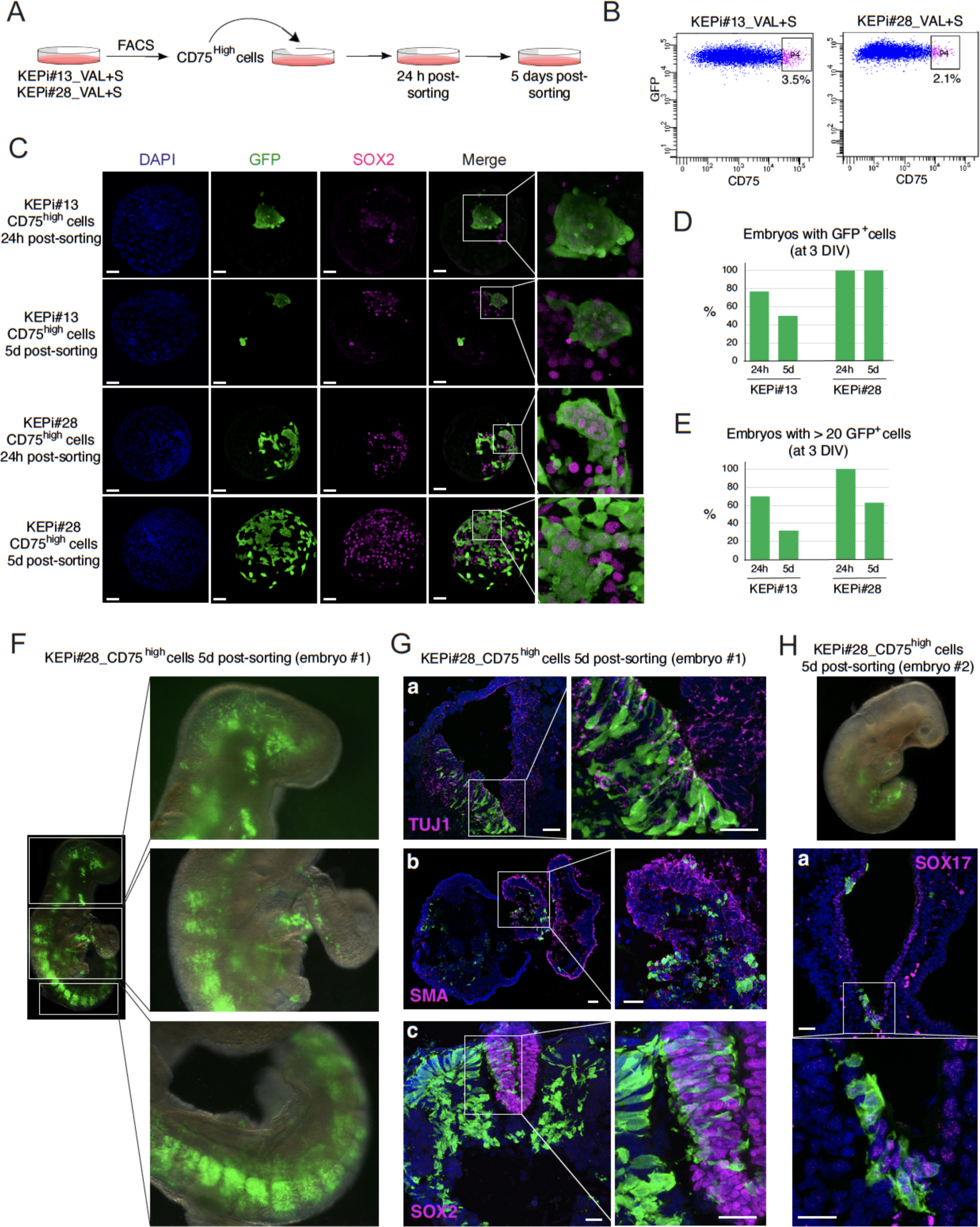
KEP_VAL cells with the highest CD75 expression exhibit massive colonization capacity. (**A**) Experimental scheme of CD75^high^ cell subpopulation analysis. (**B**) Flow cytometry analysis of KEP#13_VAL+S and KEP#28_VAL+S cells showing the FACS-sorted cells. (**C**) Confocal images of late-blastocyst-stage rabbit embryos (3DIV) acquired after microinjection of KEPi#13_CD75^high^ and KEPi#28_ CD75^high^ cells into morula-stage (E2) embryos (scale bars: 50 µm). (**D**) Percentage of embryos with GFP^+^ cells with each cell line. (**E**) Percentage of embryos with > 20 GFP^+^ cells with each cell line. (**F**) Chimeric embryo collected at E10.5. (**G**) Confocal images of sections from a E10.5 chimeric embryo (#1) showing GFP^+^ cells in the neural tube (co-labeled with TUJ1 and SOX2), in the heart (co-labeled with SMA), and in the somites. **(H**) Confocal images of sections from a E10.5 chimeric embryos (#2) showing GFP^+^ cells in the gut epithelium (co-labeled with SOX17). (F-H) Scale bars: 20 µm.

To investigate the ability of CD75^+^ cells to contribute to germ layer differentiation, we injected a total of 192 morulae with KEPi#28_CD75^high^ cells. These were cultured for 24 hours before being transferred to surrogate female rabbits. Of the 39 embryos collected at E10, five (12%) showed extensive integration of iPSC-derived GFP^+^ cells (**Figs. 7F, S7B**). Phenotypic analysis of GFP^+^ cells in embryo sections revealed their differentiation into several cell types: SOX2^+^ neuronal progenitors and TUJ1^+^ neurons, SMA^+^ (smooth muscle actin) cardiac cells, and SOX17^+^ gut cells (**Fig. 7G**). A notable contribution of the iPS cells was observed in the head mesenchyme and the somitic mesoderm along the anterior-posterior axis. GFP^+^ cells were also present in the amnion, surface ectoderm (LAMININ^+^ cells), and gut epithelium (SOX17^+^ cells (**Fig. 7H**). Overall, these results demonstrate the high efficiency of KEPi_CD75^high^ cells in contributing to the development of tissues from ectodermal, mesodermal, and endodermal origins in rabbit foetuses.

## Discussion

In this study, we identified three genes, *KLF2*, *PRMT6*, and *ERAS*, that enable rabbit iPSCs to acquire molecular and functional features of the naïve state. These features include long-term self-renewal in a KOSR+FGF (KF)-free culture medium supplemented with LIF, activin A, PKC and WNT inhibitors, a transcriptional shift reflective of early blastocyst, the reactivation of the 2^nd^ X-chromosome, and the capacity to efficiently colonize rabbit preimplantation embryos. *KLF2* encodes a transcription factor participating in the maintenance of the naïve state of pluripotency in mice (Jiang et al. 2008; Hall et al. 2009; Yeo et al. 2014; Qiu et al. 2015; Jeon et al. 2016; Dunn et al. 2018; Yamane et al. 2018). KLF2 can also promote the primed to naïve state conversion in human PSCs (Hanna et al. 2010; Takashima et al. 2014). *Prmt6* encodes an arginine methyltransferase that influences mESC self-renewal by controlling H3R2me levels. Specifically, knockdown of *Prmt6* leads to downregulation of pluripotency genes and induction of expression of differentiation markers (Lee et al. 2012). Eras, a member of the RAS family, is known to enhance the growth and tumorigenicity of mouse ESCs. However, Eras expression is not essential to mouse ESC pluripotency (Takahashi et al. 2003). In our study, we showed that overexpression of KLF2 was necessary and sufficient for iPSC self-renewal in VALGöX, but iPSCs underwent a more comprehensive conversion when both PRMT6 and ERAS were included in the reprogramming cocktail. Specifically, overexpression of the three transgenes resulted in the expansion of a cell subpopulation expressing higher levels of the naïve marker CD75 and devoid of H3K27me3 foci, indicating exceptionally immature cells. These immature cells were not observed in KEP_KF cell populations, and were barely detectable in KE_VAL, and KP_VAL cells. They emerged when KEP cells are derived in VALGöX culture conditions. Similarly, embryo colonization ability was virtually zero for parental iPSCs cultivated in KF, low for KEP-KF cells, moderate to high for KEP cells cultivated in VALGöX, and very high for the CD75^high^ sub-population. Overall, these results indicate a synergy between the VALGöX culture regimen and KEP transgenes to enable rabbit iPSCs to enter a functional naive state including an embryo colonization capability. Interestingly, a fraction of the chimeric embryos exhibited KEP-derived cells in both the epiblast and trophoblast, suggesting that some KEP cells have the characteristics of expanded pluripotency as defined in mice, pigs, and humans (Yang et al. 2017a; Yang et al. 2017b; Gao et al. 2019).

It remains unclear how PRMT6 and ERAS work together with KLF2 to accomplish this conversion. PRMT6 is known to exert various effects in cancer cells, influencing processes such as cell growth, migration, invasion, and apoptosis. PRMT6 upregulation has been linked to global DNA hypomethylation by impairing the chromatin binding of UHRF1, a co-factor of DNMT1, which results in passive DNA demethylation (Veland et al. 2017). PRMT6 has also been shown to have a carcinogenic effect by activating the AKT/mTOR pathway in endometrial cancer and prostate cancer cells (Almeida-Rios et al. 2016; Jiang et al. 2020; Chen et al. 2022). In various cancer cell lines, including breast, lung, colorectal, and cervical, as well as in osteosarcomas, PRMT6 was found to inhibit the expression of cyclin-dependent kinase inhibitors (CKI) p21cip1/waf1, p27kip1, and p18ink4c, or to mitigate the association of p16ink4a with CDK4. This results in accelerated cell cycles and the inhibition of cellular senescence (Kleinschmidt et al. 2012; Wang et al. 2012; Bao et al. 2019; Tang et al. 2020). In breast and prostate cancer cell lines, PRMT6 was found associated with regulation of motility and invasion through upregulation of thrombospondin-1 (TSP-1) and downregulation of matrix metalloproteases (Kim et al. 2013). Additionally, in breast cancer cells, PRMT6 was found to suppress IGFBP3 expression, thereby reducing apoptosis (Kim et al. 2010). Indeed, the diverse roles of PRMT6 in cell cycle, migration, and apoptosis can be seen as integral to the primed-to-naïve state conversion and embryo colonization capacity of KEP cells. These functions could potentially enhance the cells’ fitness, improve their resistance to harsh environments, and boost their adaptability. This interplay of mechanisms suggests that PRMT6 might be a key factor in facilitating more efficient chimera generation, but further research is needed to understand the intricate dynamics at play. The role of ERAS in enhancing the transition from a primed-to-naïve state and fostering subsequent chimeric competence is more elusive. Similar to PRMT6, ERAS is known to activate the AKT pathway and accelerate the mitotic cycle in mouse ESCs (Takahashi et al. 2003; Takahashi et al. 2005). We can conjecture that its exogenous overexpression could have a substantial impact on the growth of rabbit iPSCs, which might explain why ERAS has consistently emerged in our screens. However, further investigation is necessary to confirm this hypothesis and comprehend the role of ERAS in the context of embryo colonization.

The cells KEPc and KEPi provide a unique opportunity to identify transcriptomic alterations and cellular functions that distinguish cells with high aptitude for embryonic colonization from those with little to no competence. Our findings reveal that chimeric competence is strongly associated with the transcriptional repression of genes involved in MAPK, WNT, HIPPO, and EPH signaling pathways. These findings corroborate previous studies which underscored the role of MAPK, WNT, HIPPO pathway inhibition in capturing and preserving naïve pluripotency (Qin et al. 2016; Weinberger et al. 2016; Shi et al. 2020; Bayerl et al. 2021). The down-regulation of EPH receptors and ephrin ligands is a new finding. It is consistent with the overarching role of EPH-ephrin signaling in counteracting self-renewal, whether in progenitor cells of the nervous system, skin and intestinal stem cells, or in cancer stem cells (Kania and Klein 2016; Arora et al. 2023). Further investigation is necessary to clarify the role of EPH-ephrin signaling in naïve pluripotency and embryo colonization capability. Furthermore, the chimeric competence of KEPc and KEPi cells is associated with the transcriptional activation of genes involved in amino-acid metabolism, NF-kB signaling, and p53 pathway. These observations align with prior findings made in naïve *versus* primed human PSCs that underscored the KEGG pathway *valine, leucine, and isoleucine biosynthesis* as well as the GO terms *signal transduction in response to DNA damage*, *intrinsic apoptotic pathway in response to DNA damage,* and *signal transduction by p53 class mediator* as naïve-specific terms (Ernst et al. 2015). These pathways might have a viral role in maintaining genomic integrity in response to an increased energy demand.

KEPi cell populations contain a subset expressing the naïve marker CD75 at much higher levels than average. Furthermore, these CD75-high cells exhibit remarkable embryonic colonization capacity: nearly all pre-implantation embryos are colonized by day 3 in vitro (3DIV). Additionally, some of these colonized embryos can produce chimeric fetuses, in which the contribution from PSC cells is significant, affecting derivatives of ectoderm (neural tube, cell surface ectoderm), mesoderm (somitic mesoderm, heart), and endoderm (gut). These findings demonstrate our success in generating rabbit PSC cells with the ability to produce systemic chimeras comparable to mouse PSCs.

## Materials and Methods

### Media composition, culture, and electroporation

Mouse embryonic fibroblasts (MEFs) were prepared from 12.5-day-old embryos of OF1 or DR4 mice (Charles River). Conventional rabbit iPSCs B19, NaiveRep_KF, and KEP_KF cell lines were routinely cultured on mitomycin C-treated MEFs (1.6 × 10^4^ MEFs/cm^2^) in medium designated as KF comprising Dulbeccos’s Modified Eagle Medium (DMEM)/F12 supplemented with 1% non-essential amino acids, 1% solution of 10,000 U/ml penicillin, 10,000 U/ml streptomycin, 2 mM L-glutamine, 1 mM sodium pyruvate, 100 μM 2-β-mercaptoethanol, 20% knockout serum replacement (KOSR), and 10 ng/ml FGF2. The culture medium was replaced every day, and cells were routinely dissociated every 2 or 3 days into single cells by treatment with 0.05% trypsin–EDTA. In NaiveRep_KSR/LIF cells, FGF2 was replaced by LIF (medium designated as KSR/LIF). In NaiveRep_FCS/LIF cells, KSR and FGF2 were replaced by FCS and LIF (medium designated as FCS/LIF). NaiveRep_VAL and KEP_VAL cells were cultured on Matrigel in a medium designated as VALGöX, comprising MEF-conditioned N2B27 basal media supplemented with 1% non-essential amino acids, 1% solution of 10,000 U/ml penicillin, 10,000 U/ml streptomycin, 29.2 mg/ml L-glutamine, 1 mM sodium pyruvate, 100 μM 2-β-mercaptoethanol, 50 µg/ml activin A, 10,000 U/ML leukemia inhibitory factor (LIF), 250 µM Vitamin C, 2.5 µM Gö6983, and 2.5 µM XAV939. The culture medium was replaced every day, and cells were routinely dissociated every 2 or 3 days into single cells by treatment with 0.05% trypsin–EDTA. Shield1 (10 to 1000 nM) was added as indicated.

To establish the NaiveRep cell line, 10^6^ B19_GFP cells were co-transfected using Lipofectamine 2000 (Invitrogen) with 1 μg of *pCAG-Cas9D10A-sgDPPA2(x2)* plasmid and 1 μg of *pMA-5’DPPA2-mKO2-PGK-neo-pA-3’DPPA2* template. G418 (250 μg/ml) was applied for 7 days prior to colony picking and DNA analysis. “On target” integration of the DNA template was verified by PCR using the following primer pair: *5’-GCAGAGTAAGCCCACTCCAG-3’* and *5’-GACCATCGGCAGGAAAGTTA-3’.* Heterozygote integrations were distinguished from homozygotes integrations using the following primer pairs: *5’-GCAGAGTAAGCCCACTCCAG-3’* and *5’-ACGAGAAAAGCAA GCAGGTC-3’*. In a second step, 10^6^ B19_GFP_DPPA2^mKO2^ cells were co-transfected using Lipofectamine 2000 (Invitrogen) with 1 μg of *pEOS^tagBFP^*plasmid plus 2 μg of the PBase-expressing vector *pCAGPBase* (Wang et al. 2008). Puromycin (1 μg/ml) was applied for 7 days to select stable transfectants.

To establish PiggyBac (PB) transgenic lines, 10^6^ cells were co-transfected using Lipofectamine 2000 (Invitrogen) with 1 μg of PB plasmid plus 2 μg of the PBase-expressing vector *pCAGPBase* (Wang et al. 2008). Stable transfectants were selected in G418 (250 μg/ml), hygromycin (200 μg/ml), or puromycin (1 μg/ml) for 7 days.

### Virus production, infection of rabbit iPSCs, and cDNA library screening

293T cells were transfected with a DNA mixture containing 10 µg of the *pGRev* plasmid encoding the vesicular stomatitis virus glycoprotein envelope, 10 µg of *pSIV3+* plasmid encoding the gag, pol, tat, and rev proteins, and 13 µg of the *pW10* plasmid carrying the lentiviral genome (Negre et al. 2000) using the calcium phosphate precipitation technique. The following day, cells were incubated with 5 ml of fresh DMEM and further cultured for 24 h. The supernatant was then collected, filtered (0.45µm), cleared by centrifugation (3000 rpm, 15 min), and concentrated by ultracentrifugation (90,000 g, 90 min).

Prior to infection, 2.4 x 10^5^ NaiveRep_KF were dissociated with Trypsin-EDTA, and cells were transferred to fresh medium containing viruses at MOI’s of 10 to 50, in the presence of 6 µg/ml polybrene. Cells were incubated in suspension for 5 h at 37°C before being re-plated on fresh feeder cells in KF, FCS/LIF, or KSR/LIF culture media, or on Matrigel-coated dishes in VALGöX medium, at a density of 100 cells/cm^2^. Colonies showing mKO2 and/or tagBFP fluorescence were picked after 7 to 10 days and expanded.

### Genomic PCR amplification of proviral DNAs

Proviral DNAs were identified by PCR on genomic DNA, either using the Quick-Load Taq2X Master Mix enzyme (NEB, M0271S), or with the Q5 Hot Start High-Fidelity DNA polymerase (NEB, M0493S) and the addition of GC enhancer to aid in the amplification of GC-rich segments. The primer sequences are given in **Table S2**.

### Cell microinjection, embryo culture, and embryo transfer to surrogate rabbits

All procedures in rabbits were approved by the French ethics committee CELYNE (approval number APAFIS#6438 and APAFIS #39573. Rabbit embryos were produced by ovarian stimulation. Sexually mature New Zealand white rabbits were injected with follicle-stimulating hormone and gonadotropin-releasing hormone, followed by artificial insemination or breeding, as previously described (Teixeira et al. 2018). Eight-cell-stage embryos (E1.5) were flushed from explanted oviducts 36–40 h after insemination and cultured in a 1:1:1 mixture of RPMI 1640 medium, DMEM, and Ham’s F10 (RDH medium; Thermo Fisher Scientific) at 38°C in 5% CO_2_ until cell microinjection. Eight cells were microinjected inside early morula (E2.8) stage rabbit embryos. The embryos were further cultured for 24h in a 1:1 mixture of cell culture medium and RDH (with or without Shield1 as indicated). Embryos were then either further cultured in RDH medium (with or without Shield1) after removal of the mucus coat with pronase, or transferred into surrogate mothers. On the day of transfer, eight embryos were transferred to each oviduct of the recipient by laparoscopy (Besenfelder et al. 1998). Seven days after transfer, post-implantation embryos (E10.5) were recovered by dissection of the explanted uterine horns.

### Immunofluorescence analysis of cells and embryos

Epifluorescent and phase contrast imaging of live cells and chimeric embryos, following injection with GFP^+^ cells (at the blastocyst stage, 3DIV, or at E10.5 upon collection), was conducted using a conventional fluorescence microscope (TiS; Nikon). This microscope is equipped with DAPI (Ex 377/50, Em 447/60), mKO2 (Ex 546/10, Em 585/40) and GFP (Ex 472/30, Em 520/35) filters. The images were analyzed using NIS-Elements imaging software. For immunofluorescence analysis, cells and preimplantation embryos were fixed in 4% PFA for 20 min at room temperature. After three washes in phosphate-buffered saline (PBS), they were permeabilized in PBS-0.5% TritonX100 for 30 min and blocked in 2% BSA for 1 h at room temperature. For 5’methylcytosine immunolabeling, cells were permeabilized in PBS-0.5% Triton for 15 min, washed in PBS for 20 min, then incubated in 2M HCl for 30 min prior blocking as above. In all cases, cells and embryos were subsequently incubated with primary antibodies diluted in blocking solution overnight at 4°C. Primary antibodies include: anti-SOX2 (Bio-Techne, ref AF2018, dilution 1:100), anti-GFP (Invitrogen, ref A10262, dilution 1:200), anti-OCT4 (StemAB, ref 09-0023, dilution 1:200), anti-DPPA5 (R&D systems, ref AF3125, dilution 1:100), anti-OOEP (Abcam ref 185478, dilution 1:100), anti-ubiquityl-Histone H2A Lys119 (Cell Signaling, ref #8240, dilution 1:300), anti-Tri-methyl Histone H3 Lys27 (Cell Signaling, ref #9733, dilution 1:400), anti-Tri-methyl Histone H3 Lys4 (Cell Signaling, ref #9751, dilution 1:400), anti-Histone H3 (acetyl K14) (Abcam, ref ab52946, dilution 1:400), anti-Histone H3 tri methyl K9 (Abcam, ref ab8898, dilution 1:400), anti-Histone H3 (asymmetric di methyl R2) (Abcam, ref ab175007, dilution 1:400), anti-5-methylcytosine (EMD Millipore, ref MABE146, dilution 1:400), and anti-CD75 (Abcam, ref ab 77676, dilution 1:1000). After two washes (2 x 15 min) in PBS, they were incubated in secondary antibodies diluted in blocking solution at a dilution of 1:400 for 1 h at room temperature. Primary antibodies (diluted 1:500 for all of them) include: donkey anti-goat (Alexa Fluor 555) (Invitrogen ref A21432), donkey anti-chicken (Alexa Fluor 488) (Jakson ImmunoResearch, ref 703-545-155), donkey anti-mouse (Alexa Fluor 647, Invitrogen, ref A21448), donkey anti-rabbit (Alexa Fluor 555, Invitrogen, ref A31572) and donkey anti-rabbit (Alexa Fluor 647)(Invitrogen, ref A31573). Finally, they were transferred through several washes of PBS before staining DNA with DAPI (0.5μg/mL) for 10 min at room temperature. Cells and embryos were analyzed by confocal imaging (DM 6000 CS SP5; Leica). Z-stacks were acquired with a frame size of 1024 x 1024, a pixel depth of 8 bits, and a z-distance of 1µm between optical sections for cells and 5µm for embryos. Acquisitions were performed using an oil immersion objective (40x/1.25 0.75, PL APO HCX; Leica) for cells and a water immersion objective (25×/1.25 0.75, PL APO HCX; Leica) for embryos.

Post-implantation embryos (E10.5) were fixed in 2% paraformaldehyde (PFA) prior to inclusion in OCT and cryosectioned. Transversal sections, 20 μm thick, were prepared and mounted on Superfrost Plus glass slides (Thermo Scientific) before being stored at −20°C. For immunostaining, cryosections were air-dried for 30 min and rehydrated in phosphate-buffered saline (PBS) for 30 min. The slides were then rinsed three times in 0.5% TritonX100 in PBS and blocked in PBS supplemented with 2% bovine serum albumin (BSA) for 1 h at room temperature. Primary antibodies were incubated overnight at 4°C in 2% BSA. These included: anti-SOX2 (Bio-Techne, ref AF2018, 1:100), anti-GFP (Invitrogen, ref A10262, dilution 1:200), anti-TUJ1 (Sigma, ref T8660, 1:10000), anti-smooth muscle actin (SMA; Millipore, ref CBL171, 1:500), anti-LAMININ (Sigma, ref I9393, 1:100), and anti-SOX17 (Bio-Techne, ref AF1924, 1:25). After two washes in PBS, relevant secondary antibodies were incubated in 2% BSA for 1 h at room temperature, at 1:400 dilutions for all: Alexa Fluor 488 goat anti-chicken (Abcam, ab63507), Alexa Fluor 647 donkey anti-mouse IgG (Invitrogen, ref A21448), Alexa Fluor 647 donkey anti-goat IgG (Invitrogen, A21447), and Alexa Fluor 647 donkey anti-rabbit IgG (Invitrogen, ref A31573). Nuclear staining was performed using 4′,6-diamidino-2-phenylindole (DAPI; 0.5μg/mL in PBS) for 10 min at room temperature. Mounting was done in Vectashield with DAPI (Vectorlabs, H-1200-10). Confocal examination of the fluorescent labeling was conducted on a LEICA DM 6000 CS SP5 equipped with an Argon laser (488 nm), a HeNe laser (633 nm), and a diode at 405 nm. Acquisitions were made using oil immersion objectives (x20 and x40) with the LAS AF software (Leica). Z-stacks were acquired with a frame size of 2048 x 2048 and a z-distance of 1 µm between optical sections.

### Flow cytometry analysis and sorting

Cells were dissociated to single cell suspension with 0.1% trypsin, and incubated in 10% fetal bovine serum for 15 min at room temperature for epitope blocking. Cells were subsequently incubated with CD75 primary antibody (Abcam, ab77676; dilution 1:1000) for 30 min at room temperature, followed by goat anti-mouse IgG secondary antibody coupled to Alexa Fluor™ Plus 647 (Invitrogen, A32728; dilution 1:200) for 30 min. After several washes in PBS, cells were analyzed using LSRFortessa™ X-20 Cell Analyzer (Beckton Dickinson), and the data were analyzed using FlowJo software. The CD75^high^ sub-population was sorted using FACSAria™ III Sorter (Beckton Dickinson).

### Western Blotting

Cells were lysed in cold RIPA lysis buffer (0.5% NP-40, 1% Triton-X, 10% glycerol, 20 mM Hepes pH 7.4, 100 mM NaCl, 1 mM sodium orthovanadate, 0.1% DTT, and protease inhibitors (Roche, #05 892 970 001) for four hours at 4°C. Lysates were cleared by centrifugation for 15 min and stored at −80°C. Protein concentrations were measured using Bradford assay. For SDS-PAGE electrophoresis, 30µg of total proteins were loaded onto each well of Mini-PROTEAN TGX Stain-Free Precast Gels (10%, Biorad, # 4568031), and migrated for 45 min at 120 Volts. Precision Plus Protein Dual Color Standards (Biorad, # 1610374) were used as protein ladder. After electroporation, proteins were transferred onto membranes using Trans-Blot^®^ Turbo^TM^ RTA Midi 0.2 µm Nitrocellulose Transfer Kit (Biorad, #1704271). The membranes were subsequently blocked in TBST solution (200 mM Tris-HCl, 1.5 M NaCl, 0.1% Tween-20, 5% milk) for one hour at room temperature, prior to incubation with primary antibodies diluted in TBST at 4°C for 12 hours. Primary antibodies include: anti-V5 Tag (Invitrogen, ref R96025, 1:500, anti-HA (Sigma-Aldrich, ref H6908, 1:500), anti-Akt (pan) (Cell Signaling, ref #4691, 1:1000), anti-Phospho-Akt (Ser473) (Cell Signaling, ref #4060, 1:1000), and anti-beta-actin (Sigma-Aldrich, ref A3854, 1:10000). Membranes were incubated with HRP-conjugated secondary antibody (Jackon ImmunoResearch anti-mouse ref 211-032-171 and anti-goat ref 115-035-146, dilution 1:5000) for one h at room temperature. After serial washing in TBST, HRP activity was revealed using Clarity^TM^ Western ECL substrate (Biorad, #170-5060) and ChemiDoc^TM^ MP imaging system (Biorad).

### RNA extraction and RNA sequencing

Total RNA was isolated using trizol/chloroform protocol followed by RNeasy mini kit (Qiagen #74106) with a DNase I (Qiagen #79254) treatment. Three nanograms of total RNA were used for amplification using the SMART-Seq V4 Ultra Low Input RNA kit (Clontech) according to the manufacturer’s recommendations (10 PCR cycles were performed). cDNA quality was assessed on an Agilent Bioanalyzer 2100, using an Agilent High Sensitivity DNA Kit. Libraries were prepared from 0.15 ng cDNA using the Nextera XT Illumina library preparation kit. Libraries were pooled in equimolar proportions and sequenced (Paired-end 50–34 bp) on an Illumina NextSeq500 instrument, using a NextSeq 500 High Output 75 cycles kit. Demultiplexing was performed using bcl2fastq2 (version 2.18.12) and adapters were trimmed with Cutadapt (version 1.15 and 3.2) so that only reads longer than 10 bp were kept. Number of reads ranged from 10 to 200 million post adapter trimming. Reads were mapped to the rabbit genome (*OryCun2_ensembl92*) using tophat2. 49.8 to 58.4% (depending on samples) of the pair fragments could be uniquely mapped to gene reference.

### Image analysis

Fluorescent profiles measurement were generated using Fiji (Schindelin et al. 2012), and 3D reconstructions were performed using Imaris Viewer. Quantitative image analysis was conducted using Fiji. Briefly, after z-projection of the stack (sum slices function), nuclei were first segmented using an automatically set threshold and watershed separation. Signals to be quantified were then segmented with the masks defined by the nuclei. For segmenting the spot corresponding to the X-chromosome, the size and the circularity of the objects to be detected within each nucleus were determined with the control group. The same parameters were then used on the other groups to determine how many spots could be detected per cell. For all detected objects, the area and mean intensity were calculated and exported for statistical analysis in Microsoft Excel file format.

### Bioinformatics Analysis

Each sample was analyzed in three biological replicates produced at least one week apart. For bulk RNA-seq, count tables were generated using FeatureCounts (version 1.5.0-p2). All analyses were executed using R software (version 4.1.2). Data normalization, gene expression levels, and PCA were performed using the DESeq2 R package (version 1.34.0) (Love et al. 2014). Mean expression levels and standard deviations were calculated using the R base package (version 4.1.2) for each sample. After log2 transformation, normalized counts were used to produce heatmaps with the pheatmap R package (version 1.9.12). DESeq2 computed log2 transformed Fold Change (FC) and p-values for each feature. Differentially expressed genes (DEGs) were determined by Over-Representation Analysis (ORA) using DESeq2 results and the following thresholds: FC > 2 ∪ FC < (−2), p-value < 0.01 or < 0.05 as indicated. Venn diagrams were produced using the VennDiagram package (version 1.7.3) and volcano plots were designed using the ggplot2 package (version 3.4.1). Finally, KEGG (Kyoto Encyclopedia of Genes and Genomes) pathway enrichment analysis was performed using the EnrichR (Chen et al. 2013) web tool and the human KEGG pathways database (release 2021). Gene expression analysis at the single-cell level in rabbit embryos was carried out using data from Bouchereau et al., 2022 (GSE180048). The dataset was processed using the Seurat R package (version 4.3.0). Bulk RNA-seq data from rabbit embryos were retrieved from Bouchereau et al., 2022 ((Bouchereau et al. 2022)PRJNA743177) and were analyzed using the DESeq2 R package.

### Quantification and statistical analysis

The Mann-Whitney U test, suitable for non-normal distribution was conducted using the R base package (version 4.1.2) to compare medians between samples. A difference was considered significant if the p-value associated with the difference between medians was less than 0.05. For immunofluorescence analysis, either conventional T-test or Welch’s unequal variances t-test was employed. The SuperPlotsOfData tool (Postma and Goedhart 2019) was utilized to account for all measurements and biological replicates in these comparisons.

## Data and code availability

RNA-seq data have been deposited at GEO (GSE250288) and are publicly available as of the date of publication. To review GEO accession GSE250288, go to: https://www.ncbi.nlm.nih.gov/geo/query/acc.cgi?acc=GSE250288 Enter token qxkjqsgirlgvfod into the box Original western blot images have been deposited at Mendeley: https://data.mendeley.com/preview/fr4j7thttr?a=eb7adef1-6551-48ba-99b5-fda7d1f7e1a2.

## Acknowledgements

This work has benefited from the facilities and expertise of the high throughput sequencing core facility of I2BC (Yan Jaszczyszyn, Magali Perrois, Centre de Recherche de Gif, http://www.i2bc.paris-saclay.fr/), the cytometry core facility of CRCL (Priscillia Battiston-Montagne, Thibault Andrieux, Centre de Recherche en Cancérologie de Lyon, https://www.crcl.fr/en/platforms/flow-cytometry-core-facility/), and the staff members of the animal facility of SBRI (Stem cell and Brain Research Institute, https://sbri.fr/). This work was supported by the Agence Nationale pour la Recherche (contracts ANR-18-CE13-023, ORYCTOCELL; ANR-21-CE20-0018-01, CHROMNESS), the Fondation pour la Recherche Médicale (DEQ20170336757 to P.S.), the Infrastructures Nationales en Biologie et Santé (ANR-11-INBS-0009, INGESTEM; CRB-Anim, ANR-11-INBS-0003), the IHU-B CESAME (ANR-10-IBHU-003), the LabExs (ANR-10-LABX-73, REVIVE; ANR-10-LABX-0061, DEVweCAN; ANR-11-LABX-0042, CORTEX), and the University of Lyon within the program “Investissements d’Avenir” (ANR-11-IDEX-0007).

## Author contributions

Investigation: F.P., Y.P., H-T.P., N.D., A.M., S.R-G., M.A., N.B.

Formal analysis: Y.P., L.J., V.D.

Writing original draft, Funding acquisition: P.S., NB

Conceptualization, Supervision, Validation, Visualization, Project administration, Manuscript review and editing: P.S., M.A., N.B.

Resources: S. R-G., T.J., B.P.

